# Mechanism of DNA unwinding by hexameric MCM8-9 in complex with HROB

**DOI:** 10.1101/2023.06.12.544631

**Authors:** Ananya Acharya, Hélène Bret, Jen-Wei Huang, Martin Mütze, Martin Göse, Vera Kissling, Ralf Seidel, Alberto Ciccia, Raphaël Guérois, Petr Cejka

**Affiliations:** Institute for Research in Biomedicine, Università della Svizzera italiana (USI), Faculty of Biomedical Sciences, Bellinzona, 6500, Switzerland; Department of Biology, Institute of Biochemistry, Eidgenössische Technische Hochschule (ETH), Zürich, 8093, Switzerland; Institute for Integrative Biology of the Cell (I2BC), Commissariat à l’Energie Atomique, Centre National de la Recherche Scientifique, Université Paris-Sud, Université Paris-Saclay, Gif-sur-Yvette, 91190, France; Department of Genetics and Development, Institute for Cancer Genetics, Herbert Irving Comprehensive Cancer Center, Columbia University Irving Medical Center, New York, NY, USA; Peter Debye Institute for Soft Matter Physics, Universität Leipzig, Leipzig, 04103, Germany

## Abstract

The human MCM8-9 helicase functions in concert with HROB in the context of homologous recombination, but its precise function is unknown. To gain insights into how HROB regulates MCM8-9, we first used molecular modeling and biochemistry to define their interaction interface. We show that HROB makes important contacts with both MCM8 and MCM9 subunits, which directly promotes its DNA-dependent ATPase and helicase activities. MCM8-9-HROB preferentially binds and unwinds branched DNA structures, and single-molecule experiments reveal a low DNA unwinding processivity. MCM8-9 unwinds DNA as a hexameric complex that assembles from dimers on DNA in the presence of ATP, which is prerequisite for its helicase function. The hexamer formation thus involves two repeating protein-protein interfaces forming between the alternating MCM8 and MCM9 subunits. One of these interfaces is rather stable and forms an obligate heterodimer, while the other interface is labile and mediates the assembly of the hexamer on DNA, independently of HROB. The ATPase site composed of the subunits forming the labile interface disproportionally contributes to DNA unwinding. HROB does not affect the MCM8-9 ring formation, but promotes DNA unwinding downstream by possibly coordinating ATP hydrolysis with structural transitions accompanying translocation of MCM8-9 on DNA.

## Introduction

DNA helicases are motor proteins that move directionally along a nucleic acid phosphodiester backbone, separating two strands of a DNA double-helix. The minichromosome maintenance (MCM) proteins are a subfamily of hexameric DNA helicases belonging to the AAA+ ATPase family that function in diverse cellular processes including DNA replication and repair^1–3^. In humans, the MCM helicase family contains eight members (MCM2-9)^4^. The best characterized are the MCM2-7 proteins that form the motor of the CMG (Cdc45-MCM-GINS) DNA replicative helicase. Recent advances using biochemical and structural approaches uncovered that MCM2-7 is recruited as a single hexameric open ring together with Cdt1 by the Origin Recognition Complex (ORC) and Cdc6 to dsDNA, followed by another MCM2-7-Cdt1 complex to form an inactive double hexamer^5–8^. Subsequently, the MCM2-7 complexes are turned into two active single hexameric CMG helicases encircling single-stranded DNA (ssDNA), where MCM2-7 is stimulated by GINS and Cdc45^7, 9,10^. The loading and activation of MCM2-7 is quite complex, likely reflecting the necessity to restrict the initiation of DNA replication to take place only once during the cell cycle at the onset of S-phase.

Other AAA+ helicases, including archaeal replicative MCM proteins, also form hexamers. However, they assemble from only one polypeptide. Archaeal MCM helicases similarly form double-hexamers prior to activation, however, as all other AAA+ helicases, function as single hexameric rings during DNA unwinding^11^. The loading mechanism and activation of the MCM and other AAA+ helicases may differ, yet several characteristics are common to most members of this family characterized to date^2, 3^. These proteins contain an N-terminal oligonucleotide-oligosaccharide (OB-fold) domain, which is involved in protein oligomerization and DNA binding. The conserved C-terminal parts of the protein harbor the AAA+ ATPase. Characteristically, a functional ATPase site is reconstituted from two adjacent subunits: the Walker A and B motifs are provided by one polypeptide, while another component, termed the arginine finger, belongs to the neighboring subunit^12–14^. Consequently, ATP is hydrolyzed at the interface of the two subunits, typically in a sequential manner along the ring structure, and hexamer formation is hence a prerequisite for DNA unwinding activity. While the MCM complexes may exist as planar rings, during DNA unwinding they form a spiral staircase-like structure that employs a "hand-over-hand" mechanism to move along ssDNA^2, 3^.

The MCM8 and MCM9 proteins are only found in multicellular eukaryotes^4, 15^. They are missing in fungi and nematodes, but present in most plants and vertebrates, while *Drosophila* only contains an MCM8 homologue^16–18^. The MCM8 and MCM9 proteins associate together (MCM8-9) and are similarly thought to form hexamers^19–21^, yet the oligomeric state of the full-length human proteins remains uncharacterized. While MCM8 has the standard domain structure, MCM9 contains an unusual C-terminal extension downstream of the AAA+ ATPase site^22^. In humans, defects associated with MCM8-9 were linked to primary ovarian failure and hence infertility^23–26^, and defects or overexpression of MCM8-9 may promote cancer^27–31^. Most reports to date suggest that MCM8-9 functions in meiosis and in DNA repair, particularly in homologous recombination (HR) in response to DNA interstrand crosslinks (ICLs), as well as to maintain replication fork stability^17, 18, 22, 32–36^. Mutant cell lines show only minor sensitivity to ionizing radiation or bleomycin, suggesting that MCM8-9 is not a universal DNA break repair factor^37, 38^. Rather, MCM8-9 may function in recombination assisting ICL repair in the context of stalled replication forks, possibly alongside Fanconi anemia proteins^37–39^. MCM8-9 was also proposed to support recombination-based DNA synthesis to allow the extension of the invading DNA strand^37, 38^. However, MCM8-9 was also suggested to function at the onset of recombination during DNA end resection in conjunction with the MRN complex^40^ or to help facilitate RAD51 loading^41^. Therefore, there are reports that implicate MCM8-9 to act both upstream and downstream of RAD51 during HR. Beyond recombination, it was proposed that MCM8-9 may unwind DNA during mismatch repair^42^, an observation that was supported by the identification of mutant alleles associated with the Lynch syndrome or microsatellite instability^35, 43, 44^. Finally, while MCM8-9 is not required for general DNA replication^45^, it may have a residual function in DNA synthesis in the absence of MCM2-7, particularly on damaged DNA templates^46^. The functions of MCM8-9 to date have been inferred from cellular experiments. Due to its multiple roles associated with pleiotropic phenotypes upon depletion, it is apparent that the absence of biochemical insights makes the assignment of a precise molecular role for MCM8-9 difficult.

Recently, several groups identified a protein named HROB (homologous recombination factor with OB-fold, also termed MCM8IP or C17orf53)^37, 39, 47^. Defects in HROB cause pronounced meiotic defects, ICL sensitivity and recombination impairment that resembled and were epistatic with defects in MCM8-9^37, 39, 47^. RAD51 loading in HROB-deficient cells was normal, leading to persistent RAD51 foci, hinting that HROB with MCM8-9 might act at the postsynaptic HR stage downstream from RAD51^37^. While HROB has no apparent catalytic activity, it was found to physically interact with MCM8-9, and to stimulate its DNA unwinding activity^39^. The siRNA-mediated depletion of HROB compromised the accumulation of MCM8 at DNA repair foci but not *vice versa*, suggesting that HROB acts upstream of MCM8-9^37^. It was inferred that HROB may help load MCM8-9 on DNA, yet the mechanism of MCM8-9 loading on DNA and activation was not demonstrated^37, 39, 47^.

Using molecular modeling, ensemble and single molecule biochemistry, as well as cell biology, we define here the physical and functional interactions between HROB and MCM8-9. HROB interacts with both MCM8 and MCM9 subunits. The OB-fold domain of HROB does not support DNA binding, but it is essential for its interaction with MCM9 and hence the stimulation of the MCM8-9 helicase activity. MCM8-9 in conjunction with HROB prefers to bind and unwind branched DNA structures, and single-molecule experiments with magnetic tweezers revealed that DNA unwinding by the ensemble is not processive. We show that MCM8-9 forms hexamers that assemble from dimers on DNA, and ATP helps to lock the MCM8-9 ring on single-stranded DNA (ssDNA). Unexpectedly, HROB does not affect the oligomerization, loading or closing of the MCM8-9 ring on ssDNA, and does not affect its substrate preference. Rather, the reconstituted assays demonstrate that HROB primarily promotes MCM8-9 downstream of its assembly on DNA to stimulate specifically translocation and productive DNA unwinding, possibly by coordinating structural transitions of the MCM8-9 ring during DNA unwinding.

## Results

### HROB interacts with both subunits of the MCM8-9 heterodimer

Previously, HROB was found to interact with MCM8-9^37, 39, 47^. In particular, we showed that HROB bound MCM8 via a region spanning residues 396 to 413 of HROB^39^. The primary structure of HROB contains a proline-rich region (PRR18) with an unknown function, and an oligonucleotide-oligosaccharide-binding (OB-fold) domain (Fig. 1a). Overall, HROB is predicted to be largely unstructured, containing 79% disordered regions according to MobiDB^48^, with the exception of the OB-fold domain. OB-folds often mediate protein-DNA and/or protein-protein interactions, and the role of the OB-fold in HROB remained unknown. To determine its function, we expressed and purified an internally-truncated HROB variant lacking the OB-fold domain spanning residues 492-575, HROB-ΔOB (Fig. 1a,b). HROB-ΔOB was indistinguishable from the wild type protein in ssDNA binding (Fig. 1c and Extended Data Fig. 1a), but entirely deficient in promoting the helicase activity of MCM8-9 (Fig. 1d,e). These data show that during DNA unwinding, the OB-fold domain of HROB is not involved in DNA binding, and may be primarily responsible for the interaction with MCM8-9, representing a yet uncharacterized interaction interface. The function of the DNA binding domain in HROB remains unknown.

**Fig. 1.**
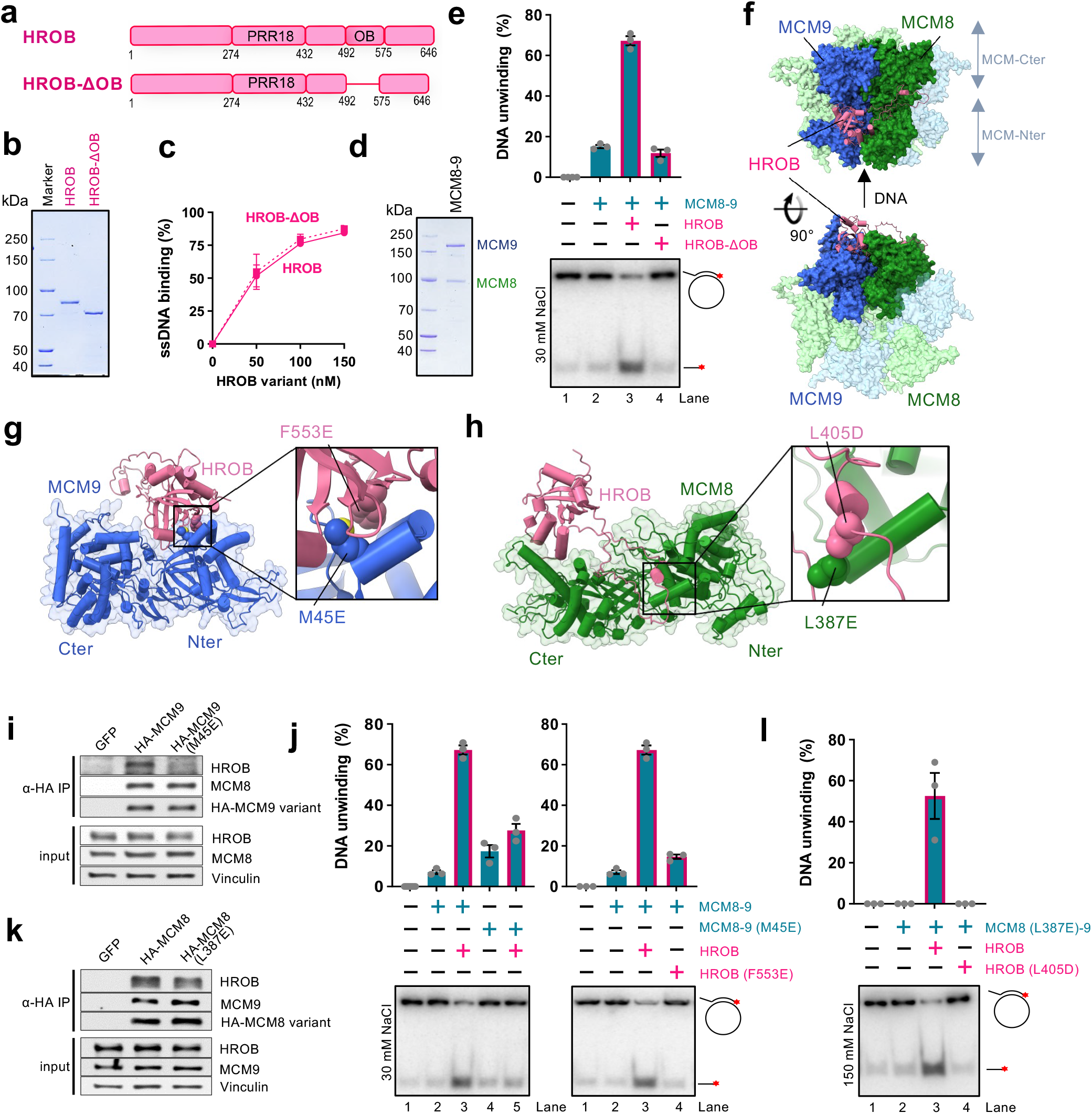
HROB interacts with both subunits of the MCM8-9 heterodimer. a. A schematic representation of HROB and HROB-ΔOB, an internal deletion mutant of HROB lacking the OB-fold domain. b. Purified wild type human FLAG-HROB and FLAG-HROB-ΔOB used in this study. c. Quantification of assays such as shown in panel Extended Data Fig. 1a (dashed: HROB-ΔOB, solid: HROB). Error bars, SEM; n = 3. d. Purified wild type human MBP-MCM9 and FLAG-MCM8 (MCM8-9 complex) used in this study. e. DNA unwinding by MCM8-9 (100 nM) and either HROB or HROB-ΔOB (both 50 nM), using M13-based circular DNA substrate with 30 mM NaCl. The red asterisk indicates the position of the radioactive label. Top, quantitation; error bars, SEM; n = 3; bottom, a representative experiment. f. A schematic representation of human HROB binding to human MCM8-9 hexameric complex modelled using AlphaFold2. g. A schematic representation of the interaction between human MCM9 and HROB. Highlighted are the residues HROB-F553 and MCM9-M45 that interact with each other. F553E and M45E point mutations in HROB and MCM9, respectively, were tested in further assays. h. A schematic representation of the interaction between human MCM8 and HROB. Highlighted are the residues HROB-L405 and MCM8-L387 that interact with each other. The L405D and L387E point mutations were tested in further assays. i. MCM9-M45E mutation disrupts MCM9-HROB interaction in cell extracts. Lysates from HEK293T cells expressing GFP, HA-MCM9-WT or HA-MCM9-M45E were subjected to HA-immunoprecipitation. Immunoblotting of HROB, MCM8 and HA are presented along with a loading control, vinculin. Shown is a representative of two independent experiments. j. DNA unwinding with indicated mutants (100 nM) to test the impact of disrupted HROB (50 nM) interaction with MCM9 using M13-based circular DNA substrate with 30 mM NaCl. HROB wild type with MCM8-9 wild type is replotted as in panel e for reference. The red asterisk indicates the position of the radioactive label. Top, quantitation; error bars, SEM; n = 3; bottom, a representative experiment. k. MCM8-L387E mutation disrupts MCM8-HROB interaction. Lysates from HEK293T cells expressing GFP, HA-MCM8-WT and HA-MCM8-L387E were subject to HA-immunoprecipitation. Immunoblotting of HROB, MCM9 and HA are presented along with a loading control, vinculin. Shown is a representative of two independent experiments. l. DNA unwinding with indicated mutants to test the impact of disrupted HROB (25 nM) interaction with MCM8 (50 nM) using M13-based circular DNA substrate with 150 mM NaCl. The red asterisk indicates the position of the radioactive label. Top, quantitation; error bars, SEM; n = 3; bottom, a representative experiment.

We next set out to define the physical interaction between HROB and MCM8-9. AlphaFold2 predicted that MCM8-9 forms a hexamer (Fig. 1f), in agreement with recent cryoEM data obtained with human MCM8-9 N-terminal domains^21^. The structural model indicated a possible interaction between the OB-fold domain of HROB (446-580) and the hinge between the N-and C-terminal domains of MCM9, located on the outer side of the MCM8-9 hexameric complex (Fig. 1f,g). Next, upstream of the HROB OB-fold, an unstructured region spanning residues 362-440 was predicted to bind MCM8 over an extensive surface located at the hinge between the N- and C-terminal domains of the helicase (Fig. 1f,h), in agreement with the fragment analysis reported previously^39^.

To validate the predicted structures, several mutations were designed at positions located in the interface between HROB and MCM8, as well as between HROB and MCM9, respectively (Fig. 1g,h). Two conserved and apolar residues, HROB-F553 and MCM9-M45, anchored in the interface between the OB-Fold of HROB and the N-terminal domain of MCM9 were substituted into a glutamate to destabilize their assembly (Fig. 1g and Extended Data Fig. 1b). We observed that the single point mutations either on the MCM9 side (M45E), or within the OB-fold of HROB (F553E) dramatically reduced physical interactions as assayed with recombinant proteins *in vitro* or in cell extracts (Fig. 1i and Extended Data Fig. 1c,d). Consequently, both single point mutants were strongly impaired in DNA unwinding, confirming the importance of the interaction interface between MCM9 and HROB (Fig. 1j). To assess the importance of the interaction of HROB with the MCM8 subunit, we focused on the most conserved positions in the disordered tail of HROB and analyzed their contacts with MCM8 residues. HROB-L405 was modelled to form an apolar contact with MCM8-L387 (Fig. 1h). This contact is predicted by AlphaFold2 to be conserved in non-vertebrate and plant species (Extended Data Fig. 1e,f). Probing for this second interacting region, mutating individually L387E in MCM8 or L405D in HROB reduced the physical interaction, although to a lesser extent than the mutations targeting the HROB-MCM9 interface (Fig. 1k and Extended Data Fig. 1c,d). In DNA unwinding, the individual mutations of the HROB-MCM8 interface initially did not cause notable defects (Extended Data Fig. 1g). However, when MCM8-L387E and HROB-L405D mutations were combined and more restrictive reaction conditions were employed, we observed a notable impairment of DNA unwinding (Fig. 1l). Together, we establish that there are at least two functionally important interfaces between HROB and the MCM8-9 complex, one on each subunit of the complex.

### Inhibition of MCM8-9 by CDK phosphorylation

The MCM9 protein was fused with an MBP-tag at the N-terminus, while MCM8 contained an N-terminal FLAG-tag to facilitate expression and purification^39^. We noted that the presence of the MBP tag did not affect DNA unwinding by MCM8-9 with HROB (Extended Data Fig. 1h). As MCM8-9 was unstable during lengthy purification and the tag cleavage, the MBP tag was retained for most experiments, unless noted otherwise. Cyclin-dependent kinase (CDK) regulates the activity of MCM2-7 with respect to the cell cycle stage on multiple levels^49, 50^. We have noted in human cell extracts that the MCM9 subunit exhibited changes in electrophoretic mobility upon treatment with λ-phosphatase, indicating that MCM8-9 may be also subject to phosphorylation (Fig. 2a). Interestingly, the C-terminal extension of MCM9 contains a large number of putative CDK phosphorylation sites with unknown function (Extended Data Fig. 2a). To test whether phosphorylation of MCM8-9 regulates its capacity to unwind DNA *in vitro*, we prepared the recombinant MCM8-9 complex in insect cells without or with phosphatase inhibitors, and treated the complex with λ-phosphatase (for the one prepared without phosphatase inhibitors) during purification. We noted that recombinant MCM9 exhibited changes in its electrophoretic mobility, resembling what we found in human cell extracts (Extended Data Fig. 2b). DNA binding and unwinding by MCM8-9 was generally inhibited by salt (Extended Data Fig. 2c,d). We observed that under physiological ionic strength conditions (150 mM salt), phosphatase treatment of phosphorylated MCM8-9 stimulated DNA unwinding (Fig. 2b), in agreement with the comparison of non-phosphorylated and phosphorylated MCM8-9 variants expressed in insect cells, in assays at 150 mM NaCl (Extended Data Fig. 2c). The moderate inhibition of the MCM8-9 complex upon phosphorylation could be recapitulated upon treatment with recombinant CDK, showing that the observed inhibitory effects are primarily dependent on the phosphorylation of MCM8-9 CDK sites (Fig. 2c,d and Extended Data Fig. 2e). In contrast, the phosphorylation status of MCM8-9 did not affect its affinity to DNA, as measured by electrophoretic mobility shift assays (EMSA, Extended Data Fig. 2f). In addition, phosphorylation of MCM8-9 did not alter its interaction with HROB (Extended Data Fig. 2g). The regulation of MCM8-9 helicase by phosphorylation resembled that of MCM4-6-7, which was similarly found to be inhibited by CDK2-Cyclin A^50^.

**Fig. 2.**
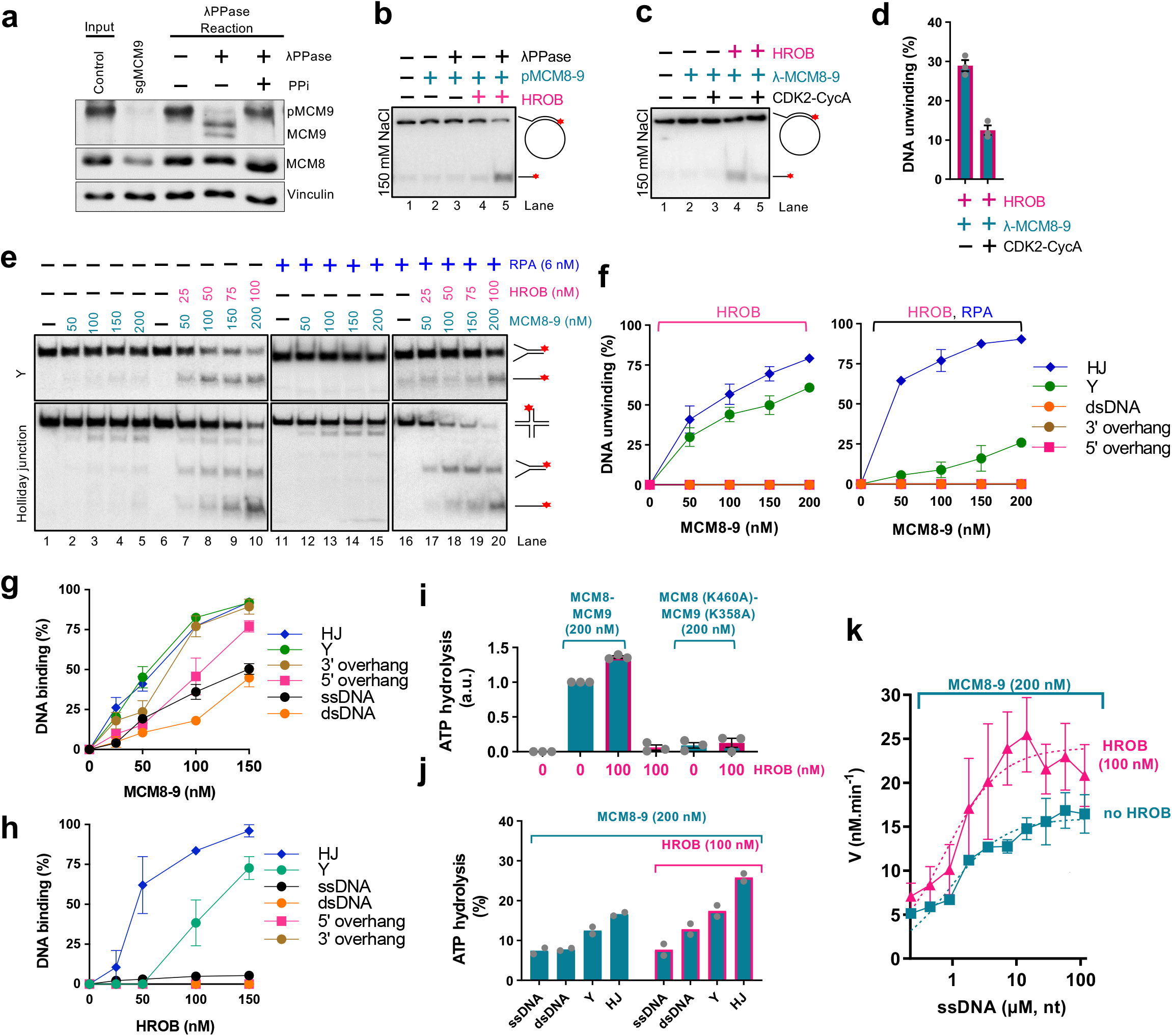
MCM8-9 with HROB unwinds branched DNA structures. a. Endogenous MCM9 is phosphorylated. Lysates from HEK293T control cells (input) were treated with lambda phosphatase (λPPase) in the absence or presence of phosphatase inhibitors (PPi) and resolved by SDS-PAGE supplemented with Phos-Bind. Immunoblotting of MCM9 and MCM8 is presented along with a loading control, vinculin. Specificity of the MCM9 band is demonstrated by the loss of signal in lysates from cells stably transduced with LentiCRISPRv2-Puro sgMCM9. Shown is a representative of two independent experiments. b. DNA unwinding by phosphorylated human pMCM8-9 (100 nM) with or without lambda phosphatase (λPPase) treatment, with or without HROB (30 nM) using M13-based circular DNA substrate with 150 mM NaCl. The red asterisk indicates the position of the radioactive label. Shown is a representative experiment. c. DNA unwinding by non-phosphorylated human MCM8-9 (100 nM), treated or not with CDK2-CycA, with or without HROB (30 nM) at 150 mM NaCl using M13-based circular DNA substrate with 150 mM NaCl. The red asterisk indicates the position of the radioactive label. Shown is a representative experiment. d. Quantification of assays such as shown in panel c. Error bars, SEM; n = 3. e. DNA unwinding by MCM8-9 without or with HROB and RPA, as indicated, with Y and Holliday junction (HJ) DNA substrates with 1 mM ATP, 5 mM magnesium acetate and 15 mM NaCl. Representative assays are shown. f. Quantification of helicase assays such as shown in panel e and Extended Data Fig. 2i. Error bars, SEM; n = 3. g. Quantification of DNA binding assays with MCM8-9 such as shown in Extended Data Fig. 2j. Error bars, SEM; n = 3. h. Quantification of DNA binding assays with HROB such as shown in Extended Data Fig. 2j. Error bars, SEM; n = 3. i. Quantification of ATP hydrolysis (expressed as arbitrary units, a.u., normalized to wild type MCM8-9 alone as 1) by 200 nM of wild type MCM8-9 and ATP-binding deficient mutant MCM8 (K460A)-MCM9 (K358A) with or without HROB. 7.2 μM (in nucleotides) M13-based circular ssDNA substrate was used as a co-factor. Error bars, SEM; n = 3. j. ATP hydrolysis (expressed in % of total ATP hydrolysedby MCM8-9 with and without 100 nM HROB in the presence of various oligonucleotide based DNA substrates (7.2 μM, in nucleotides) and RPA (0.58 μM). Bars show range; n = 2. k. Relationship between ATP hydrolysis by MCM8-9 (200 nM) and the concentration of DNA (μM, in nucleotides) without and with HROB (100 nM). V is the rate of ATP hydrolysis. Error bars, SEM; n = 4.

### MCM8-9 together with HROB unwinds branched DNA structures

To better understand the function of MCM8-9 and HROB, we set out to define its substrate preference. Starting with the circular M13-based structures^39^, we observed that MCM8-9 and HROB most efficiently displaced an oligonucleotide with a 5’ ssDNA overhang, followed by an oligonucleotide with a 3’ overhang or no overhang at all (Extended Data Fig. 2h). In fact, under relaxed experimental conditions, the 5’-tailed oligonucleotide was displaced by MCM8-9 even without HROB, although HROB promoted the unwinding of all the M13-based structures tested (Extended Data Fig. 2h). All MCM proteins unwind DNA with a 3’-5’ polarity^51^. Considering the established polarity, as well as our observation that a 5’-overhang substrate was unwound most efficiently (Extended Data Fig. 2h), suggests that while MCM8-9 translocates on the M13 DNA strand in a 3’-5’ direction, the encounter with the 5’ flap promotes strand displacement. Such preference would be consistent with the replicative MCM proteins from archaea or eukaryotes (MCM2-7); in fact, the outer *Sulfolobus solfataricus* MCM surface was proposed to dynamically interact with the 5’-tail of the strand being displaced, while the MCM ring translocates in a 3’-5’ direction on the opposite DNA strand^52^.

As the M13 ssDNA forms complex secondary structures, we next turned to oligonucleotide-based substrates to better define the MCM8-9 DNA unwinding preference. We observed that MCM8-9 and HROB unwound most efficiently a Holliday (4-way) junction (HJ) followed by a Y-structure, while 3’ and 5’ overhang as well as fully double-stranded DNA were not unwound at all under our conditions (Fig. 2e,f and Extended Data Fig. 2i). DNA unwinding of the HJ substrate was marginally stimulated by human RPA, while RPA inhibited the unwinding of the Y-structure, possibly because of competition for the single-stranded region of DNA substrate (Fig. 2e,f). DNA unwinding was strongly stimulated by HROB in all cases (Fig. 2e). The preferential unwinding of the Holliday junction substrate was unexpected, showing that a ssDNA tail is not required for the MCM8-9 helicase. The additional preference for the Y-structure was in agreement with the data obtained with the M13 substrates: we believe that MCM8-9 is loaded on the 3’-ended arm and translocates 3’-5’ towards the duplex, where it encounters the 5’ flap from the other DNA strand, which in turn stimulates DNA unwinding.

The DNA unwinding specificity of the MCM8-9 and HROB ensemble was at least in part a consequence of their DNA binding preference: MCM8-9 bound with the highest affinity Holliday junctions and Y-structured DNA (Fig. 2g and Extended Data Fig. 2j). The apparent *K*^m^, corresponding to MCM8-9 concentration when 50% DNA was bound, was ∼55 nM for HJ and Y-structured DNA. Other substrates, such as overhang DNA, were bound only slightly less. The observation that overhanged DNA is instead a poor substrate for DNA unwinding suggests that ssDNA overhangs are not sufficient to load and activate the MCM8-9 and HROB complex for productive DNA unwinding. HROB also preferentially bound HJ and Y-structured DNA, while overhanged or fully duplex DNA were almost not bound at all (Fig. 2h). We conclude that MCM8-9 is a moderately active DNA helicase that prefers to act on branched DNA. HROB stimulates the unwinding of all DNA structures tested.

### HROB promotes the ATPase activity of MCM8-9

DNA unwinding by MCM8-9 and HROB depends on ATP hydrolysis^39^. We observed that MCM8-9 exhibited ATPase activity that was strongly stimulated by DNA (Fig. 2i,j), which differs from replicative MCMs including MCM2-7 that display ATPase activity independently of DNA^12, 14, 53^. Mutation of the Walker A motif in both MCM8 (K460A) and MCM9 (K358A) subunits of the heterodimer drastically reduced the observed ATPase activity, demonstrating that it is intrinsic to the MCM8-9 complex (Fig. 2i). MCM8-9-HROB showed the highest ATPase activity with Holliday junction and Y-structured substrates, suggesting that the observed ATPase activity may reflect translocation of MCM8-9 along DNA, or DNA unwinding (Fig. 2j). We also note that ATPase activity was generally quite low.

HROB did not possess a detectable ATPase activity (Fig. 2i), as expected, and stimulated the ATPase of MCM8-9 ∼1.5-2-fold (Fig. 2i,j). The stimulatory effect of HROB on the MCM8-9 ATPase was notably less pronounced than its effect on DNA unwinding, suggesting that HROB makes the ATPase activity of MCM8-9 more productive. HROB was proposed to recruit MCM8-9 to DNA^37, 39^, and a modest stabilization of the protein-DNA species was indeed observed in the presence of HROB in electrophoretic mobility shift assays (EMSA, Extended Data Fig. 2d). However, the EMSA assays are difficult to interpret as both MCM8-9 and HROB bind DNA, and it was not clearly apparent whether DNA binding was additive or synergistic (Extended Data Fig. 2d). To learn more about the effect of HROB on MCM8-9, we next varied DNA concentration to obtain kinetic parameters for ATP hydrolysis by MCM8-9 without or with HROB (Fig. 2k). The curves clearly did not fit a Michaelis-Menten equation, indicating a complex regulation. Nevertheless, HROB stimulated the maximal rate of ATP hydrolysis (*V*_max_) ∼ 1.5-fold, while the DNA concentration that supports 1/2 of *V*_max_, corresponding to the affinity for DNA, was largely unaffected by HROB. These results suggested that HROB does not primarily recruit MCM8-9 to DNA, but may promote its activity post recruitment.

### Single-molecule experiments reveal low processivity of DNA unwinding by MCM8-9

To estimate the DNA unwinding processivity of MCM8-9 and HROB, we varied the length of the duplex DNA region annealed to M13-based ssDNA. The unwinding efficacy decreased ∼2-fold when the dsDNA region increased from 30 to 90 bp, suggesting that DNA unwinding by MCM8-9 and HROB is likely not very processive (Fig. 3a). We next set out to define the helicase activity of the MCM8-9 and HROB ensemble more quantitatively using single molecule magnetic tweezers. The approach monitors DNA unwinding as a function of a DNA length difference, such as between dsDNA and ssDNA. As opposed to ensemble gel-based techniques, the single-molecule approach allows to study individual active molecules, and it is unlikely to be biased by a proportion of potentially inactive proteins. To this point, we first employed a dsDNA substrate with a 5’-overhang at its end used for Sgs1 or yeast and human Dna2/DNA2 (Extended Data Fig. 3a)^54, 55^. We failed to detect any MCM8-9-HROB activity using this construct even at elevated concentrations or increased temperatures, while effective unwinding was observed with human BLM as a control (Extended Data Fig. 3b,c). A failure to detect DNA unwinding may reflect a low unwinding processivity of the MCM8-9-HROB complex that is below the resolution limit using this DNA construct.

**Fig. 3.**
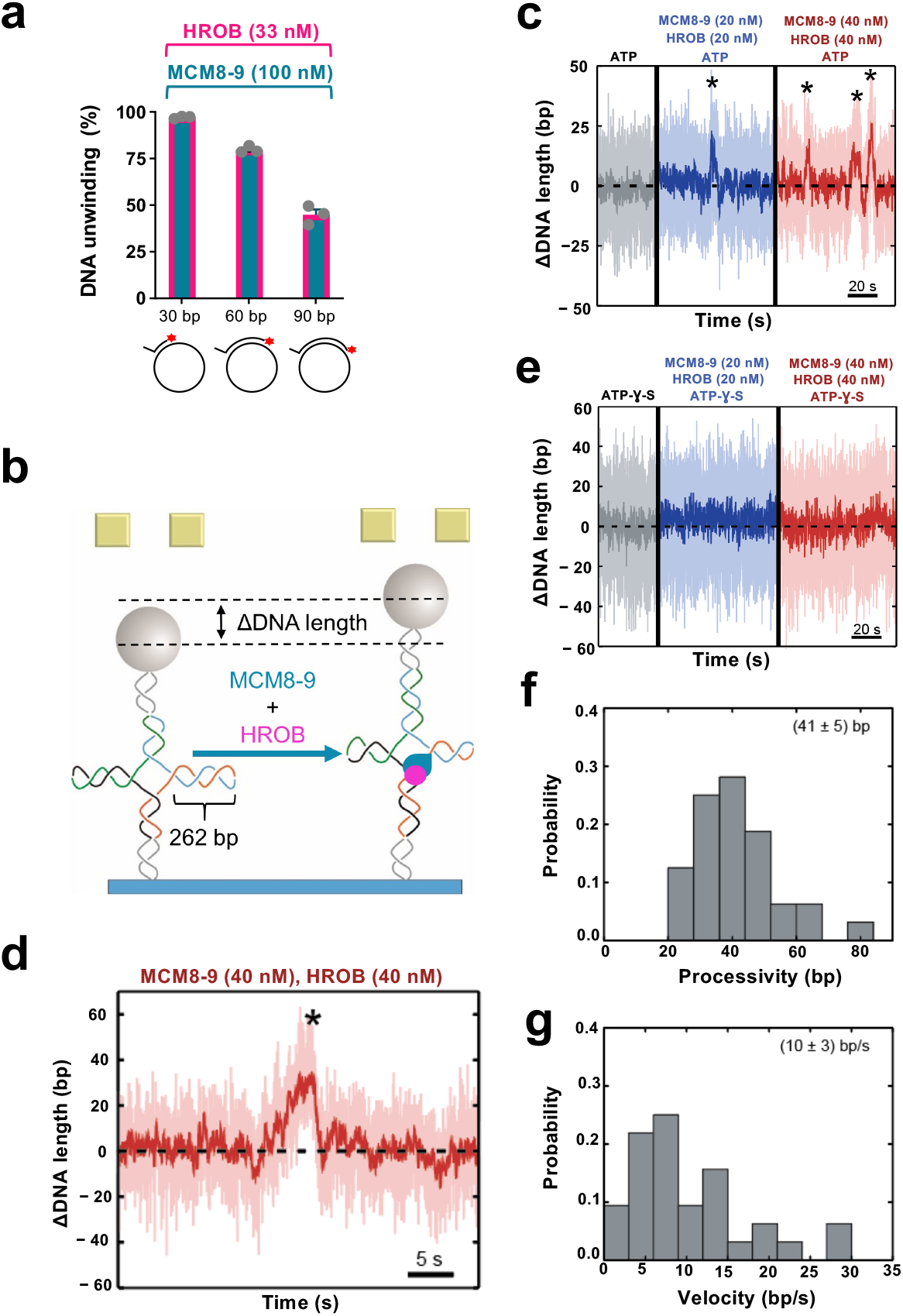
Single molecule analysis of the MCM8-9 helicase with HROB. a. Quantitation of gel-based assay showing DNA unwinding by MCM8-9 (100 nM) and HROB (33 nM), using substrates with different lengths of duplex DNA (overhang length 20 nt), as indicated with 14 mM NaCl. The red asterisk indicates the position of the radioactive label. Error bars, SEM; n = 3. b. A schematic representation of the magnetic tweezer setup and the investigated Holliday junction construct with 262 bp in each arm. When the protein ensemble is added, MCM8-9 with HROB can translocate in the direction of the arms, causing the measured DNA length to increase. c. Activity of MCM8-9 and HROB with ATP, as indicated. Successive DNA unwinding events (highlighted by *), resulting in a net-increase of DNA length, were observed. d. Magnified exemplary trace for an unwinding event by MCM8-9 and HROB. e. MCM8-9 with HROB do not unwind DNA with non-hydrolysable ATP-γ-S instead of ATP. f. Probability distribution of DNA unwinding processivity by MCM8-9 with HROB, with a mean of 41 ± 5 bp, of events leading to DNA extension, n = 32. g. Probability distribution of DNA unwinding velocity by MCM8-9 with HROB, with a mean of 10 ± 3 bp sec^-1^, of events leading to DNA extension, n = 32.

Considering that MCM8-9 efficiently unwinds HJs, we next turned to a HJ construct, with mobile arms of 262 bp in length (Fig. 3b). The unwinding or branch migration of the DNA arms would lead to a larger DNA length difference, providing greater sensitivity (Fig. 3b and Extended Data Fig. 3d). Similarly to the ensemble measurements, we did not observe any activity of MCM8-9 or HROB individually, even at increased concentrations (Extended Data Fig. 3e,f). However, the presence of both proteins in a 1:1 molar ratio resulted in short-ranged gradual unwinding events (Fig. 3c,d). No unwinding was observed in a buffer supplemented with non-hydrolysable ATP-γ-S instead of ATP, demonstrating that the detected events were caused by active motor activity rather than DNA remodeling due to protein binding (Fig. 3e). Due to the symmetric nature of the Holliday junction, two distinct event types were detected. First, we observed gradual apparent DNA lengthening, with a mean velocity of 10 ± 3 bp/sec and a mean processivity of 41 ± 5 bp (Fig. 3b,d,f,g). We also observed gradual shortening events, with a mean velocity of -(11 ± 4) bp/sec and a mean processivity of -(40 ± 7) bp (Extended Data Fig. 3d,g,h,i). The first event types originate from the MCM8-9 complex translocating from the branching point in the direction of the Holliday junction arms (Fig. 3b), whereas the second type corresponds to the complex translocating in the direction of the Holliday junction stem (Extended Data Fig. 3d). We identified ∼75% of the events corresponding to the first event type (net upwards movement), whereas the remaining 25% belonged to the second type (net downwards movement). The non-symmetric occurrence of the two events most likely originates from the impact of the externally applied force (10 pN). Nevertheless, the mean unwinding velocity and processivity was similar in both cases (Fig. 3f,g and Extended Data Fig. 3h,i), showing that the applied force likely did not affect the motor function of the helicase ensemble. We note that the DNA unwinding velocities are comparable to reports for the yeast CMG helicase that translocates on ssDNA at 5–10 bp/sec^56^ and that of 4.5 ± 1.6 bp/s for *Drosophila* CMG^57^. Our single-molecule experiments reinforce the idea that HROB stimulates the helicase function of the MCM8-9 complex and that MCM8-9-HROB *per se* is a helicase with a limited processivity, in agreement with our ensemble assays.

### HROB promotes DNA unwinding by hexameric MCM8-9

MCM8 interacts with MCM9, and the stability of the proteins in human cell extracts is partially dependent on each other^32, 33, 41^. *Drosophila* only contains MCM8, raising questions whether complexes consisting of a single MCM8 or MCM9 human polypeptides may also be functional. We observed that MCM8 and MCM9 can be expressed and purified on their own from insect cells (Fig. 4a). Individually expressed MCM8 and MCM9 were largely monomeric as determined by mass photometry (Extended Data Fig. 4a,b). However, single MCM8 or MCM9, or a combination of the individually expressed proteins did not show any DNA unwinding activity, without or with HROB, in contrast to MCM8-9 proteins expressed and purified together as a complex (Fig. 4b,c). These experiments indicated that not only both MCM8 and MCM9 are required for DNA unwinding, but also that the co-expression of both subunits is necessary, likely to achieve proper folding of the proteins (Fig. 4c). HROB is largely monomeric (Extended Data Fig. 4c). We also attempted to co-express HROB with MCM8-9, but this did not further increase the activity of the purified MCM8-9, suggesting that HROB is likely not required for the proper assembly of MCM8-9 upon expression (Extended Data Fig. 4d). Accordingly, HROB interacts with MCM8-9 rather transiently and could not be co-purified as a complex (Extended Data Fig. 4e).

**Fig. 4.**
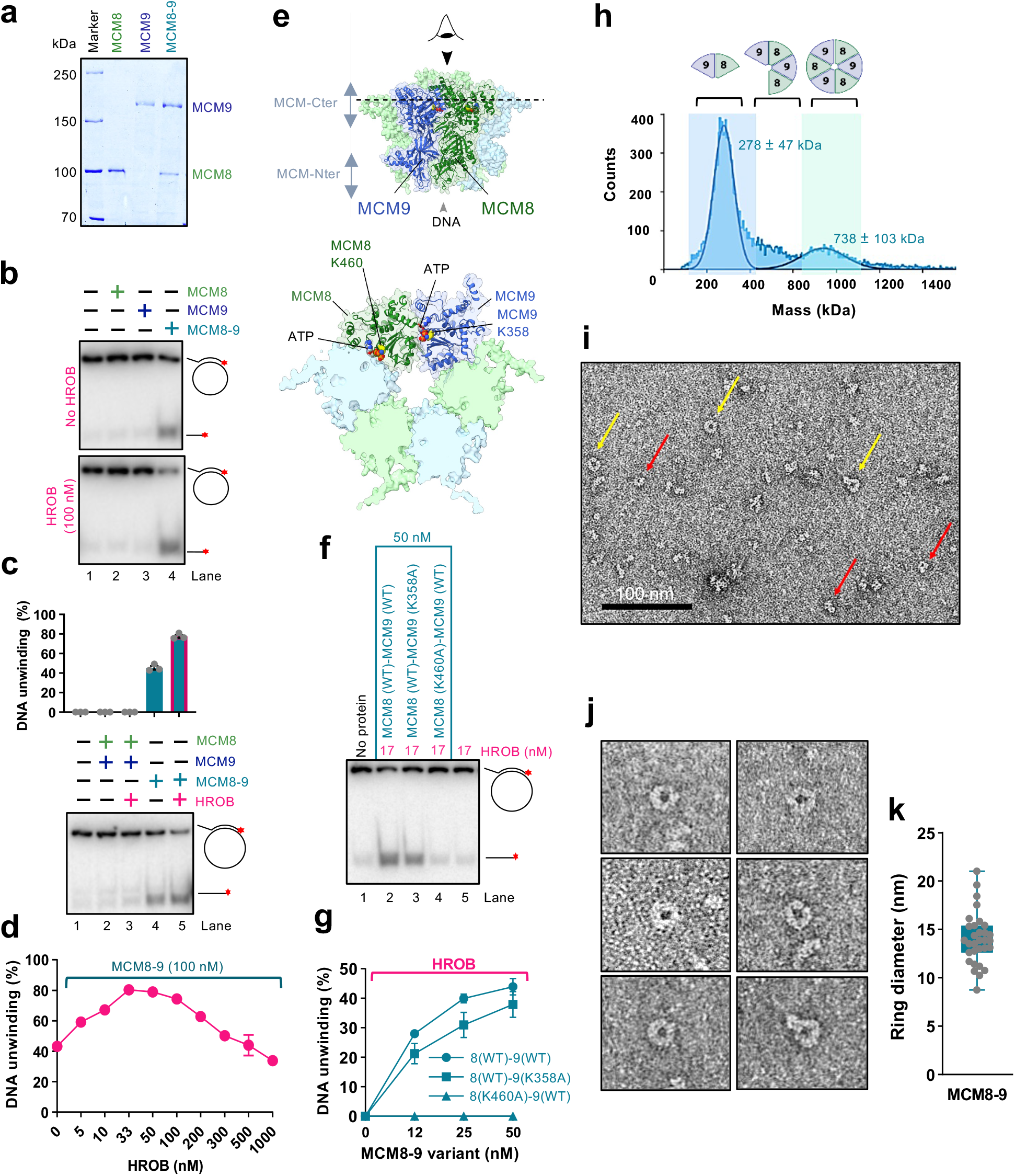
HROB promotes DNA unwinding by hexameric MCM8-9. a. Purified wild type human FLAG-MCM8, MBP-MCM9 and FLAG-MCM8-MBP-MCM9 heterodimer complex used in this study. b. MCM8, MCM9 (single polypeptides, 100 nM) and the MCM8-9 complex (co-expressed, 100 nM) were used in helicase assays with and without HROB (100 nM) using M13-based circular DNA substrate with 21 mM NaCl. The red asterisk indicates the position of the radioactive label. A representative assay is shown. c. MCM8 and MCM9, combined upon individual expression (lanes 2 and 3) or co-expressed (lanes 4 and 5), all 100 nM, were used in helicase assays with or without HROB (100 nM) using M13-based circular DNA substrate with 21 mM NaCl. The red asterisk indicates the position of the radioactive label. Top, quantification; error bars, SEM; n = 3; bottom, a representative experiment. d. Quantitation of DNA unwinding experiments by human MCM8-9 in the presence of increasing concentrations of HROB such as shown in Extended Data Fig. 4g. Error bars, SEM; n = 3. e. AlphaFold2 model depicting the positions of ATPase sites at the interfaces of MCM8 and MCM9 subunits within the hexameric complex. Walker A lysine of MCM8 (K460) and MCM9 (K358) are shown. The lower cartoon represents a view from the C-terminal end of the complex. f. DNA unwinding by wild type or ATP binding-deficient MCM8-9 variants with HROB, as indicated using M13-based circular DNA substrate with 25 mM NaCl. The red asterisk indicates the position of the radioactive label. A representative assay is shown. g. Quantification of helicase assays such as shown in panel f. HROB was used at 1/3 concentration of MCM8-9 (nM). Error bars, SEM; n = 3. h. Measured molecular weight distributions of human MCM8-9 using mass photometry. FLAG-MCM8-MBP-MCM9 was used, corresponding to a theoretical molecular weight of 266 kDa for a heterodimer and 798 kDa for a hexamer. Error, SD. i. Negative staining transmission electron micrograph of human MCM8-9 (220 nM). Yellow arrows indicate clearly distinguishable top views of MCM8-9 rings, while red arrows denote smaller, less well recognizable rings or side views of rings, which make up the majority of the visible particles. A representative image is shown. j. Representative transmission electron micrographs of MCM8-9 rings. k. Quantification of diameter sizes of distinguishable MCM8-9 rings as shown in panel j. Error bars, SD; n = 31.

We note that MCM8-9 and HROB unwound DNA in a concentration-dependent manner (Extended Data Fig. 4f). Interestingly, using a fixed concentration of MCM8-9, the highest stimulation of DNA unwinding was observed with a lower HROB concentration with respect to MCM8-9 (approximately 1:2-3 stoichiometric ratio) (Fig. 4d and Extended Data Fig. 4g). HROB did not promote DNA unwinding by the BLM helicase, but similarly became inhibitory together with BLM (Extended Data Fig. 4h). This experiment suggests that the inhibition of MCM8-9 by high HROB concentrations is non-specific, stemming likely from a competition for DNA (Fig. 2h).

We next tested the requirement of the ATPase activity of the individual MCM8 and MCM9 subunits for DNA unwinding. We observed that mutations of either the Walker A lysine residues K460A in MCM8 or K358A in MCM9 both reduced DNA unwinding of the heterocomplexes, although the contributions were different (Fig. 4e-g). No unwinding was observed with MCM8 (K460A)-MCM9 (WT) mutant, while only moderately reduced DNA unwinding was observed with the MCM8 (WT)-MCM9 (K358A) variant, showing that the MCM8 ATPase is more critical. Nevertheless, the ATPase activities of both subunits are important for DNA unwinding (Fig. 4e-g). The AAA+ helicase family ATPase site is formed at the interface of two polypeptides^12–14^, and a hexamer formation is thought to be necessary for DNA unwinding (Fig. 4e). Our observation that the integrity of both ATPase sites within the complex is important suggests that both the MCM8-MCM9 and MCM9-MCM8 interfaces are required for productive DNA helicase activities, although to different extents. Therefore, the MCM8-9 species active in DNA unwinding is of a higher order than a dimer, in agreement with other members of this helicase family and AlphaFold2 modeling (Fig. 4e and below). Mass photometry measurements of our recombinant MCM8-9 complex showed a mixed population of dimers, tetramers and hexamers (Fig. 4h). We note that the oligomeric state of MCM8-9 in solution did not change upon incubation with ATP or HROB (Extended Data Fig. 4i-k). We next performed imaging of the complexes by negative staining electron microscopy (Fig. 4i). Although we observed a variety of sizes and shapes, in accord with the mass photometer measurements, a small fraction of the protein complexes exhibited a ring-like structure with a cavity in the center (Fig. 4j,k), in accord with the AlphaFold2 model (Fig. 4e). The mean diameter of the clearly visible MCM8-9 rings in the negative staining transmission electron microscopy images was 14 nm, which is similar to the mean ring diameter measured in cryoEM images of the chicken MCM8-9 hexamer (13.2 nm)^21^, or the MCM2-7 hexamer from budding yeast (13.3 nm)^58^.

### ATP locks the MCM8-9 ring on ssDNA, irrespectively of HROB

To study the effects of MCM8-9 oligomerization on its biochemical activities, we next performed size exclusion chromatography. The elution profile showed a rather wide distribution without clearly-defined peaks, likely reflecting a dynamic nature of the MCM8-9 complexes. Nevertheless, a pool of early eluates showed a higher proportion of hexamers (hexamer-rich sample, ∼1:1 dimer:hexamer in protein counts or 1:3 in protein mass) (Fig. 5a,b), as opposed to a dimer-rich sample (∼3:1 dimer:hexamer in protein counts and 1:1 in protein mass) (Fig. 4h). Contrary to our expectations, the dimer-rich sample showed a higher apparent DNA unwinding activity than the hexamer-rich preparation together with HROB, showing that the MCM8-9 hexamers that assembled in solution without the DNA substrate may be inactive complexes (Fig. 5c and Extended Data Fig. 5a).

**Fig. 5.**
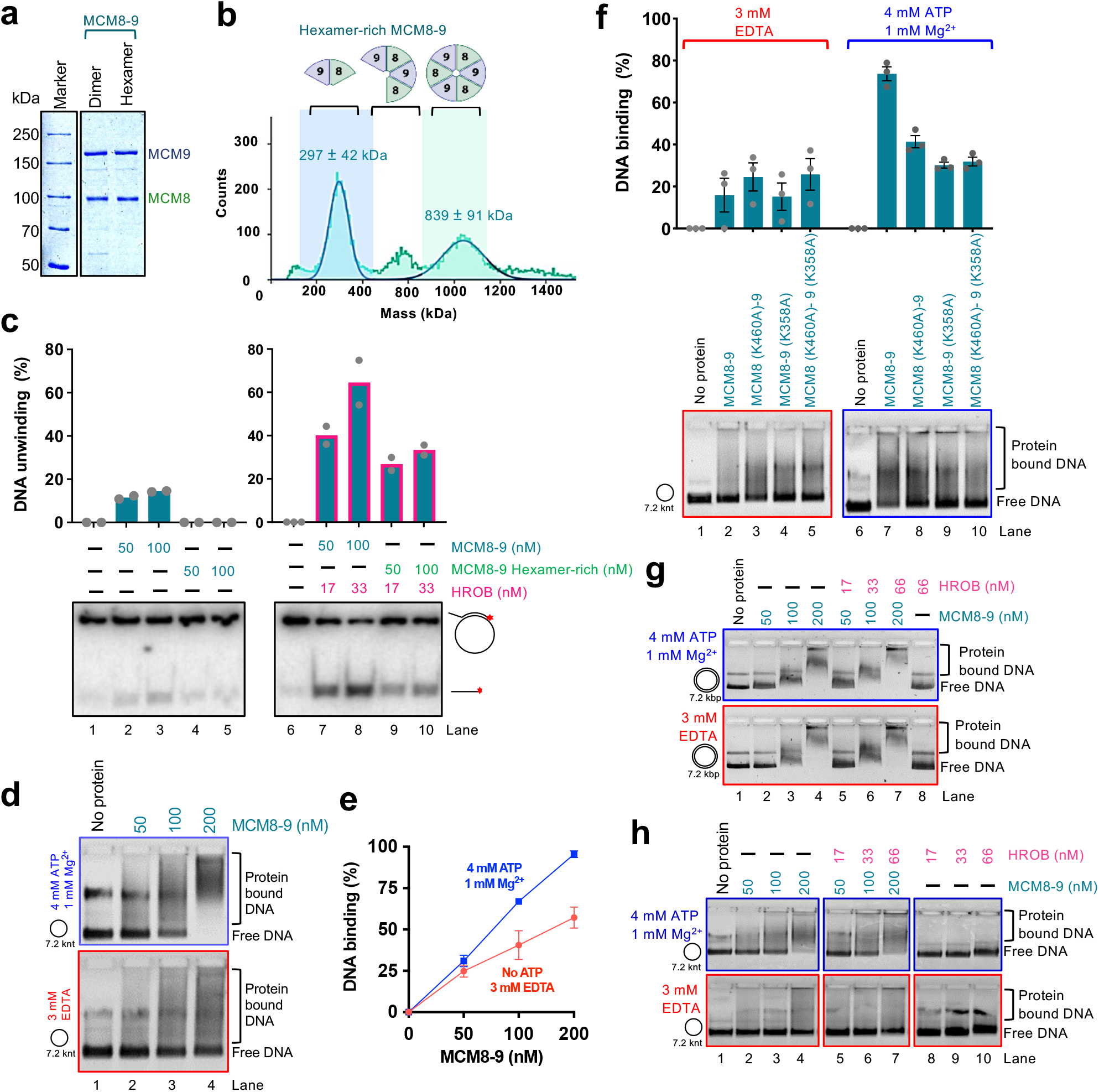
ATP locks the MCM8-9 ring on ssDNA, irrespectively of HROB. a. Purified wild type standard (dimer-rich) and hexamer-rich MCM8-9 preparations used in this study. b. Mass photometry-based molecular weight distributions of hexamer-rich MCM8-9, purified by size exclusion chromatography. Compare with Fig. 4h. Error, SD. c. DNA unwinding using standard (Fig. 4h) and hexamer-rich (Fig. 5b) preparations of MCM8-9, with and without HROB, as indicated using M13-based circular ssDNA substrate with 14 mM NaCl. The red asterisk indicates the position of the radioactive label. Top, quantification; Bars show range; n = 2; bottom, a representative experiment. d. Electrophoretic mobility shift assays with human MCM8-9, with or without ATP, as indicated, using M13-based circular ssDNA substrate. A representative experiment is shown. e. Quantification of assays such as shown in panel d. Error bars, SEM; n = 3. f. Electrophoretic mobility shift assays with ATP-binding deficient variants of human MCM8-9 (100 nM), with or without ATP, as indicated, using circular M13-based ssDNA. Top, quantification; error bars, SEM; n = 3; bottom, a representative experiment. g. Electrophoretic mobility shift assays with MCM8-9, with and without ATP, as indicated, using M13-based circular dsDNA as a substrate. Representative experiments are shown. h. Electrophoretic mobility shift assays with human MCM8-9, with and without HROB, with and without ATP, as indicated, using circular M13-based ssDNA as a substrate. Representative experiments are shown.

We next used the more active MCM8-9 dimer-rich preparation and analyzed its binding to circular and linear ssDNA, without or with ATP and without or with HROB. Several important conclusions can be made from these electrophoretic mobility shift experiments. First, MCM8-9 showed enhanced binding to circular ssDNA in the presence of ATP, as opposed to reactions without ATP (Fig. 5d,e). The enhanced binding in the presence of ATP was not observed when we used the ATP binding-deficient MCM8 (K460A)-MCM9 (K358A) mutant complex (Fig. 5f), demonstrating that the effects result from direct ATP binding to the ATPase sites of MCM8-9. These data mirror the behavior of MCM2-7, which was biochemically demonstrated to form rings on DNA^14, 59^. Next, the stabilization of DNA binding by ATP was not observed on linear ssDNA (Extended Data Fig. 5b) or on circular dsDNA (Fig. 5g). While a ring can slide off the ends of linear DNA, it remains topologically locked in place on circular DNA. These results thus suggest that ATP helps close the MCM8-9 ring around ssDNA. Finally, HROB did not affect the capacity of the MCM8-9 ring to lock onto ssDNA (Fig. 5h). We conclude that in the presence of ATP, MCM8-9 forms rings on ssDNA independently of HROB, and that the primary function of HROB is therefore downstream of MCM8-9 loading and locking onto ssDNA. Oligomerization and circularization of MCM8-9 in solution without DNA may yield inactive protein complexes.

### MCM8-9 rings assemble from heterodimers on ssDNA and form two interfaces

As described above, the MCM8-9 hexamer predicted by the AlphaFold2 model contains two distinct interfaces repeating alternatively three times along the ring structure (Fig. 6a,b, upper part). We denote the first one as interface I. When viewed from the C-terminal side of the toroid, interface I lies between subunits MCM8 and MCM9 (dashed purple line in Fig. 6a). The second, interface II, is shown by the dashed orange line in Fig. 6b between MCM9 and MCM8. HROB binds MCM8 and MCM9 across interface I (Fig. 1f). When modeling the dimer between MCM8 and MCM9, AlphaFold2 only generated interface I, suggesting that the evolutionary information encoding its structure is much stronger than that of the interface II. In each of these two interfaces, there are multiple contact points between the adjacent subunits at the N-terminal, central, and C-terminal parts of the proteins. Several mutants were designed to probe the relative importance of interfaces I and II to assess the complex formation and effects on DNA unwinding (Fig. 6a,b, lower part).

**Fig. 6.**
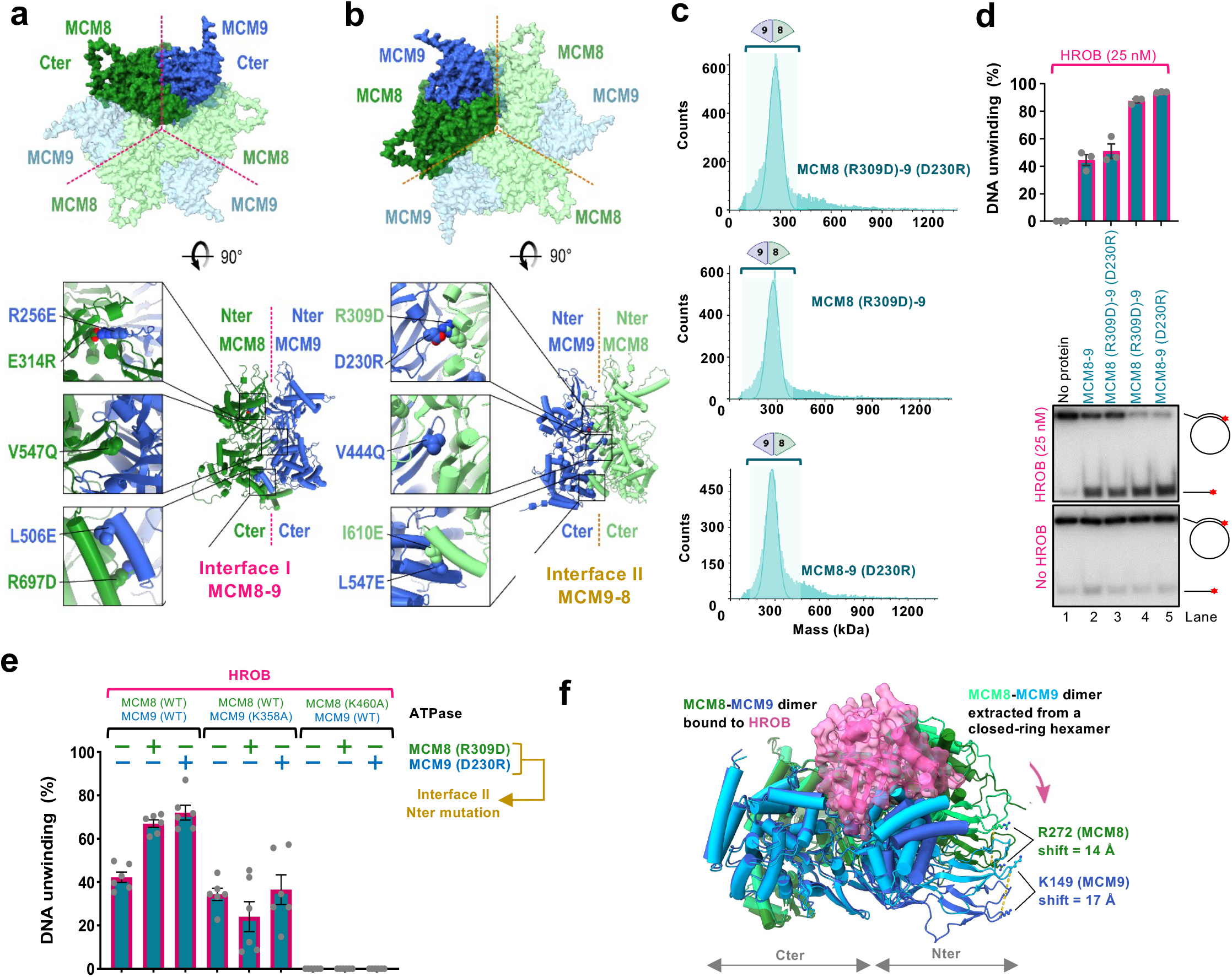
MCM8-9 rings assemble from heterodimers *via* two distinct interfaces. a. AlphaFold2 model showing a C-terminal view of the MCM8-9 hexamer (top). Interface I is indicated with a purple dotted line. Bottom part, mutations disrupting interface I were designed in the N-terminal, central, and C-terminal part of the interface. b. AlphaFold2 model showing a C-terminal view of the MCM8-9 hexamer (top). Interface II is indicated with an orange dotted line. Bottom part, mutations disrupting interface II were designed in the N-terminal, central, and C-terminal part of the interface. c. Molecular weight distributions of human MCM8-9 variants with disrupted N-terminal part of interface II measured using mass photometry. Compare with wild type MCM8-9 in Fig. 4h. d. DNA unwinding by human MCM8-9 variants with disrupted N-terminal part of interface II (50 nM), with HROB (25 nM) using circular M13-based ssDNA at 25 mM NaCl. The red asterisk indicates the position of the radioactive label. Top, quantification; error bars, SEM; n = 3; bottom, a representative experiment. e. DNA unwinding by human MCM8-9 variants (25 nM) with disrupted N-terminal part of interface II, in combination with ATPase Walker A mutations, as indicated, in the presence of HROB (8 nM) at 25 mM NaCl. The panel shows a quantification of assays such as shown in Extended Data Fig. 6c-e; error bars, SEM; n = 6. f. AlphaFold2 model depicting a structural change of the MCM8-9 dimer upon binding of HROB. For simplicity, only MCM8-9 dimer is shown, without HROB.

Interface I was mutated at five positions well exposed in the structure of each monomer (Fig. 6a). First, two mutations caused charge swapping of the salt-bridge residues at the interface between the N-terminal domains, MCM8 (E314R)-MCM9 (R256E); second, we mutated one highly conserved position in the pre-sensor-1-β-hairpin of MCM8 contacting MCM9, creating MCM8 (V547Q)-MCM9 (WT)^60^ and third, two positions in the parallel helices mediating the interaction between the C-terminal domains were replaced, creating MCM8 (R697D)-MCM9 (L506E). None of the single or double mutants produced in interface I could be purified as a heterocomplex, suggesting that the destabilization of this interface is highly detrimental to the formation of a stable dimer.

In contrast, a set of five positions mutated similarly at interface II (Fig. 6b) had no detrimental effect on the production of the MCM8-9 heterodimer and mutant proteins could be purified for biochemical analysis. The mutation located in the central pre-sensor-1-β-hairpin of MCM9, MCM9 (V444Q), structurally equivalent to MCM8 (V547Q), only partially reduced the DNA unwinding activity, suggesting that this position is less constrained than its counterpart in MCM8 (Extended Data Fig. 6a). Interface II mutants in the C-terminal domains, MCM8 (I610E)-MCM9 (L547E) were defective in DNA unwinding (Extended Data Fig. 6a). The C-terminal domains of MCM proteins harbor the ATPase/helicase domains, which likely explains the detrimental effect of these mutations.

The most interesting were mutants in interface II disrupting a salt-bridge between the N-terminal domains of MCM8 and MCM9, MCM9 (D230R) and MCM8 (R309D) (Fig. 6b). We observed that individually or combined, the mutations strongly reduced hexamer formation in solution, as apparent from mass photometry experiments (Fig. 6c), suggesting that the more labile interface II mediates the assembly of dimers into hexamers. Interestingly, while the combined MCM9 (D230R)-MCM8 (R309D) heterodimer was as active as the wild type complex in DNA unwinding (Fig. 6d and Extended Data Fig. 6a), the individual single point mutants, MCM8 (R309D)-MCM9 (WT) and MCM8 (WT)-MCM9 (D230R) showed notably elevated activity in conjunction with HROB, compared to wild type MCM8-9 (Fig. 6d). These data suggest that a moderate destabilization of the N-terminal part of interface II creates a more dynamic protein complex, which is more likely to assemble into a productive helicase on DNA, leading to overall elevated activity. Considering that the more active species MCM8 (R309D)-MCM9 (WT) and MCM8 (WT)-MCM9 (D230R) were hyperactive in helicase assays yet dimeric in solution, the data suggest that an active hexameric helicase assembles on DNA from the heterodimeric building blocks.

### Labile interface II between MCM9 and MCM8 subunits powers DNA unwinding

As introduced above, an active ATPase site of the MCM helicases is reconstituted from residues belonging to the interface of neighbor subunits^12–14^. The catalytic lysine of the Walker A motif from the MCM8 subunit K460 is predicted to lie at the labile interface II between MCM9 and MCM8, while the K358 of MCM9 maps to the stable interface I between MCM8 and MCM9 (Fig. 4e and 6a,b). As the integrity of K460 in MCM8 is more important for the helicase activity of the complex (Fig. 4f,g), the data show that ATP hydrolysis at the labile interface is more important for DNA unwinding. Therefore, although the majority of purified MCM8-9 forms stable dimers in solution mediated by the stable interface I, helicase activity is dependent on the interface II that is only assembled within the higher order complex.

To confirm the relative importance of the ATPase sites within the two interfaces, we next combined the mutations destabilizing the N-terminal part of interface II (R309D in MCM8, or D230R in MCM9) with the Walker A mutants in the C-terminal domains of MCM8 (K460A, affecting interface II) and in MCM9 (K358A, affecting interface I) (Fig. 6e). Disruption of the ATPase site in the labile interface II completely disrupted unwinding activities, while disruption of the ATPase site in the stable interface I only had a moderate effect, in all mutant combinations tested (Fig. 6e and Extended Data Fig. 6c-h). Collectively, these data support our modeling that interface I mediates stable interaction between MCM8 and MCM9, and it thus has a more structural function. Instead, the more dynamic interface II is involved in the assembly of the heterodimers into active hexameric complexes. The ATPase site at this dynamic interface contributes disproportionally more to the unwinding activity of the MCM8-9 ensemble.

## Discussion

We employed a combination of structure prediction, mutagenesis and ensemble as well as single-molecule biochemical approaches to gain insights into the mechanism of DNA unwinding by MCM8-9 and its regulation by HROB. We demonstrate that MCM8 and MCM9 proteins in solution form diverse oligomeric species ranging from dimers to hexamers, in agreement with a study in press^21^. We show that MCM8-9 hexamers are necessary for DNA unwinding in conjunction with HROB. However, hexamers assembled in solution without DNA are not active, and we provide evidence that MCM8-9 assembles on ssDNA into an active hexameric helicase from heterodimeric building blocks. HROB dramatically stimulates all DNA unwinding activities, which depends on extensive contacts with both MCM8 and MCM9 subunits of the ensemble. Interestingly, the assembly and DNA loading of MCM8-9 is not stimulated by HROB, rather, HROB preferentially stimulates DNA unwinding downstream of MCM8-9 loading and ring formation.

The ATPase sites of all MCM hexameric helicases form at the protein interfaces from residues belonging to adjacent subunits^12–14^. The MCM8-9 ring involves two distinct protein-protein interfaces, each repeating three times between the alternating MCM8 and MCM9 subunits along the ring. We find that one of these interfaces (interface I) is stable, and mediates a constitutive heterodimer formation. Mutations within this interface destabilize the dimer so that it cannot be purified. Interestingly, disruption of the ATPase site located within interface I has only a moderate effect on the DNA unwinding activity, suggesting that interface I has primarily a structural function. Interface II instead mediates the assembly of the heterodimers into hexamers. Disruption of this second interface allows the preparation of the heterodimer, and instead reduces the fraction of hexamers detected in solution. The ATPase site located at this second labile interface is instead absolutely essential for DNA unwinding.

Our conclusion that an active hexameric helicase assembles from dimers on DNA is supported by several lines of evidence. First, size exclusion chromatography-based enrichment of hexameric MCM8-9 complexes formed in solution without DNA revealed that they are largely inactive (Fig. 5a-c). Second, we find that mutations within the N-terminal region of the labile interface II reduce the fraction of hexamers in solution, but stimulate overall DNA unwinding. Third, ATP hydrolysis mediated by interface II is absolutely essential for DNA unwinding, as opposed to the ATPase site formed between the subunits of a stable dimer. Therefore, the assembly of dimers into hexamers is essential for DNA unwinding, in agreement with other members of the MCM family members^2, 3, 13^.

We find that the MCM8-9 helicase exhibits phosphorylation dependent electrophoretic mobility shift in cell extracts, and CDK phosphorylation limits its activity *in vitro* (Fig. 2a-c). MCM8-9 prefers to bind and unwind branched DNA substrates, and exhibits low DNA unwinding processivity (Fig. 2e-g, Fig. 3). HROB was initially proposed to recruit MCM8-9 to DNA, as HROB depletion disrupted the accumulation of MCM8 but not *vice versa* ^37, 39^. HROB also binds branched DNA and a weak enhancement of protein binding when both HROB and MCM8-9 were combined was observed, particularly under more restrictive conditions (Fig. 2h and Extended Data Fig. 2d). It is possible that the recruitment function is more important in cells, where MCM8-9 needs to compete with other cellular factors. Nevertheless, our data clearly demonstrate a function of HROB that is independent of MCM8-9 recruitment. We found that ATP promotes the loading and locking of the MCM8-9 ring on ssDNA, a process that is unaffected by HROB (Fig. 5d-h). This observation is in agreement with the AlphaFold2 modeling coupled with mutagenesis that positions HROB across the stable interface I through functionally important contacts with both MCM8 and MCM9 subunits (Fig. 1f-h), which provides a structural explanation why HROB does not promote hexamer formation (Extended Data Fig. 4k). We noted that AlphaFold2 model predicts a rotation of both MCM9 and MCM8 N-terminal subunits with respect to their C-terminus upon HROB binding (Fig. 6f), in agreement with a cryoEM study in press^21^. Considering two highly conserved basic residues in the N-terminal domains well positioned to bind DNA at the entrance of the helicase tore, MCM9 (K149) and MCM8 (R272), their position is predicted to shift by 17 Å and 14 Å, respectively (Fig. 6f). Such structural change may trigger activation of the helicase ensemble. We also observed that ATP hydrolysis by MCM8-9 is comparatively weakly stimulated by HROB (Fig. 2i-k), in contrast to the strong stimulation of DNA unwinding (Fig. 2e), where the stimulatory effect of HROB is much greater. Therefore, HROB facilitates productive ATP hydrolysis, possibly by coordinating the conformational changes with DNA translocation and unwinding. Additionally, a positively charged patch on HROB near the MCM8 subunit may mediate interaction with the displaced DNA strand. A surface patch of archaeal MCM was similarly proposed to explain the binding and unwinding preference for Y-shaped DNA^52^. The activation of MCM8-9 by HROB downstream of its loading may thus be reminiscent of MCM2-7 activation by Ccd45-GINS within the replicative CMG helicase complex^7, 9, 10^, or by MCM10^61, 62^.

## Methods

### Cloning, expression and purification of recombinant proteins

Human MCM8-9 and HROB were expressed in *Spodoptera frugiperda* 9 (*Sf*9) insect cells as previously described^39^, using pFB-MBP-MCM9 or pFB-FLAG-MCM8, as well as pFB-MCM8IP-FLAG. Please note that MCM8IP is the previous name for HROB^39^. The protein sequence of HROB-FLAG is shown in Supplementary Table S1. The used HROB sequence is the closest to isoform 4 (Q8N3J3-4, UniProt), because it lacks Q-484. However, the sequence also contains a common polymorphism, T-126 → P-126. We note that the isoform 1 sequence of HROB (UniProt, Q8N3J3-1) also stimulates DNA unwinding by MCM8-9, albeit to a lower extent. The MCM8-9 proteins were purified using MBP- and FLAG-tag based affinity chromatography as previously except that the MBP-tag was not cleaved^39^. Unless indicated otherwise, MCM8 and MCM9 were co-expressed and co-purified as a complex. Mutations in the expression vectors were generated using the QuikChange II XL site-directed mutagenesis kit, following manufacturer’s instructions (Agilent Technology). Sequences of all primers used for site-directed mutagenesis are listed in Supplementary Table S2. The MCM8-9 mutants were expressed and purified as the wild type complex. Wild-type HROB and mutants were purified using FLAG M2 Affinity Gel (Sigma, A2220) based column chromatography^39^.

For the expression of phosphorylated MCM8-9 (pMCM8-9) variants, *Sf*9 cells were treated with 50 nM Okadaic acid (APExBIO, A4540) to preserve proteins in their phosphorylated state. Additionally, the cell extracts were supplemented with 50 nM Okadaic acid (APExBIO), 1 mM sodium orthovanadate (Sigma), 20 mM sodium fluoride (Sigma), and 15 mM sodium pyrophosphate (Applichem) during lysis to preserve the phosphorylation status. Where indicated, pMCM8-9 was dephosphorylated with λ-phosphatase (New England Biolabs) according to the manufacturer’s instructions. For ‘‘mock’’ controls, λ-phosphatase was excluded from the reactions, and the sample was otherwise incubated as the λ-phosphatase-treated reactions. Similarly, MCM8-9 was phosphorylated by CDK2-CycA, where indicated, using standard conditions of *in vitro* phosphorylation^63^. The hexamer-rich prep of MCM8-9 was purified using the standard method for MCM8-9 purification, except by adding a size exclusion chromatography step (Superose 6 10/300 GL, GE Healthcare) in between the amylose and FLAG steps, and pooling the early eluting MCM8-9 fractions. Human RPA was expressed in *E. coli* and purified using ÄKTA pure (GE Healthcare) with HiTrap Blue HP, HiTrap desalting and HiTrap Q chromatography columns (all GE Healthcare)^64^.

### DNA substrate preparation

For DNA binding experiments with linear single-stranded DNA, oligonucleotide X12-3 HJ3 (93 nt) was labeled at the 5’ terminus with [γ-32P] (Hartmann-Analytic) and T4 polynucleotide kinase (New England Biolabs), according to standard protocols^65^. The oligonucleotide-based DNA binding, ATPase and DNA unwinding experiments that compared various DNA substrates were ssDNA (labelled oligonucleotide PC1253), dsDNA (labelled PC1253 and PC1253C), 5’ overhang (labelled PC1253 and 312), 3’ overhang (labelled PC1255 and 314), Y-structure (labelled PC1254 and PC1253) and Holliday junction (labelled PC1253 and PC1254, PC1255 and PC1256)^66^. The indicated oligonucleotides were labeled at the 3’ terminus with [α-32P] dCTP (Hartmann-Analytic) by terminal transferase (New England Biolabs) prior to annealing, according to standard procedures. Unincorporated nucleotides were removed using Micro BioSpin™ P-30 Gel Columns (Bio-Rad). Plasmid length DNA binding experiments were performed with unlabeled circular or linearized M13mp18 single (New England Biolabs, M13mp18 single-stranded DNA) or double-stranded DNA (New England Biolabs, M13mp18 RF I DNA).

A major portion of the DNA unwinding experiments were performed using 5’ overhang-M13 based circular DNA. This was prepared using oligonucleotide (M13-5’-dT 40 overhang) containing a 37 nt-long region complementary to the M13mp18(+) strand (nucleotides 6289– 6326), as well as a 40 nt-long tail at the 5’ end, which was annealed to M13mp18 ssDNA^39^. Variants of the annealed oligo with either no overhang (M13-37mer no overhang) or a 40 nt-long tail at the 3’ end (M13-3’-dT 40 overhang) were used to prepare the no overhang or 3’ overhang-M13 based circular DNA substrates, respectively. Similarly, other variants were prepared by altering the length of the annealing region to produce substrates with 30, 60 and 90 bp-long complementary region to the M13mp18(+) strand (oligonucleotides M13-5’-dT20overhang-30bp_c, M13-5’-dT20overhang-60bp_c and M13-5’-dT20overhang-90bp_c). Sequences of all oligonucleotides used for substrate preparation are listed in Supplementary Table S3. The oligonucleotides were labeled at the 3’ terminus with [α-32P] dCTP (Hartmann-Analytic) by terminal transferase (New England Biolabs) prior to annealing, according to standard procedures. Unincorporated nucleotides were removed using Micro BioSpin™ P-30 Gel Columns (Bio-Rad). The annealed substrate was purified using CHROMA SPIN™+TE-200 Columns (TaKaRa) to remove unannealed oligonucleotides.

### Electrophoretic mobility shift assays

Binding reactions (15 µl volume) were carried out in 25 mM Tris acetate pH 7.5, 3 mM EDTA, 1 mM dithiothreitol (DTT), 100 µg/ml bovine serum albumin (BSA, New England Biolabs), and DNA substrate (1 nM, in molecules). 3 mM EDTA was replaced with 4 mM ATP (Sigma, A7699) and 1 mM magnesium acetate where indicated. Proteins were added and incubated for 15 minutes on ice. Loading dye (5 µl; 50% glycerol [w/vol] and bromophenol blue) was added to the reactions and the products were separated on 4% polyacrylamide gels (ratio acrylamide:bisacrylamide 19:1, Bio-Rad) in TAE (40 mM Tris, 20 mM acetic acid and 1 mM EDTA) buffer at 4 °C. The gels were dried on 17 CHR papers (Whatman), exposed to a storage phosphor screen (GE Healthcare) and scanned by a Typhoon Phosphor Imager (FLA9500, GE Healthcare). Signals were quantified using ImageJ and plotted with GraphPad Prism. DNA binding experiments using plasmid-length substrates were performed under the same conditions except with 100 ng DNA per reaction and were run on 0.8% agarose gel at 4 °C and post-stained with GelRed (Biotium).

### Helicase assays

Helicase assays (15 µl volume) were performed in a reaction buffer (25 mM HEPES-NaOH pH 7.5, 1 mM magnesium acetate, 4 mM ATP, 1 mM DTT, 0.1 mg/ml BSA) with DNA substrate (0.1 nM, in molecules). Some reactions contained 1 mM ATP and 5 mM magnesium acetate, as indicated. Recombinant proteins were added as indicated. The reactions were incubated at 37 °C for 30 minutes and stopped by adding 5 µl 2% stop buffer (2% sodium dodecyl sulfate [SDS] [w/vol], 150 mM EDTA, 30% glycerol [w/vol] and bromophenol blue) and 1 µl of proteinase K (14–22 mg/ml, Roche) and incubating at 37 °C for 10 minutes. To avoid re-annealing of the substrate, the stop solution was supplemented with a 200-fold excess of the unlabeled oligonucleotide with the same sequence as the ^32^P-labeled one. The products were separated by 10% polyacrylamide gel electrophoresis in TBE (89 mM Tris, 89 mM boric acid, 2 mM EDTA) buffer, dried on 17 CHR paper and analyzed as described above.

### ATPase assays

ATPase assays with MCM8-9 and HROB were performed in 25 mM HEPES-NaOH pH 7.5, 5 mM magnesium acetate, 1 mM DTT, 0.1 mg/ml BSA, 0.1 mM ATP, 1 nM of [γ-^32^P] ATP (Hartmann-Analytic) with DNA as a co-factor (1 µM, in nucleotides, where indicated) in 10 µl reaction volume. Recombinant proteins were added as indicated. The samples were mixed on ice and incubated at 37 °C for 2 hours. Reactions were stopped with 2 µl of 0.5 M EDTA and separated using thin layer chromatography plates (Merck) with 0.3 M LiCl and 0.3 M formic acid as the mobile phase. Dried plates were exposed to storage phosphor screens (GE Healthcare) and scanned by a Typhoon FLA 9500 phosphorimager (GE Healthcare). Signals were quantified using ImageJ software. Spontaneous ATP hydrolysis signal from no protein lanes was removed as a background during quantitation. The percentage of ATP hydrolysis was obtained as a normalized value and expressed in arbitrary units, or as a rate of ATP hydrolysis.

### Mass photometry characterization of protein complexes

Mass photometry measurements were performed on a 2MP-0132 mass photometer (Refeyn Ltd). For the measurements, coverslips (No. 1.5 H thickness, 24 × 50 mm, VWR) were cleaned by sequential dipping in Milli-Q-water, isopropanol and Milli-Q-water followed by drying under a stream of gaseous nitrogen. Subsequently, silicone gaskets (CultureWell^TM^ Reusable Gasket, Grace Bio-Labs) were placed on the cleaned coverslips to create wells for sample loading. For mass measurements, gaskets were filled with 18 µl equilibration buffer (50 mM Tris–HCl, pH 7.5 and 100 mM NaCl) to allow focusing the microscope onto the coverslip surface. Subsequently, 2 µl protein solution was added into the 18 µl droplets and mixed to obtain 50-200 nM final protein concentration. Sample binding to the coverslip surface was monitored by recording a movie for 1 minute using AcquireMP (Refeyn Ltd) software. Data analysis was performed using DiscoverMP (Refeyn Ltd). To convert the measured optical reflection-interference contrast into a molecular mass, a known protein size marker (NativeMark^TM^ Unstained Protein Standard, Invitrogen) was measured.

### *In vitro* interaction studies

To study the interaction between recombinant MCM8-9 and HROB, *Sf*9 cells were co-infected with MBP-MCM9 and FLAG-MCM8 baculoviruses. Cells were lysed and MCM8-9 was immobilized on amylose resin and washed with wash buffer (50 mM Tris–HCl, pH 7.5, 2 mM β-mercaptoethanol, 300 mM NaCl, 0.1% (v/v) NP40, 1 mM PMSF). Resin-bound MCM8-9 was then incubated with 1 µg of HROB wild type or mutants, diluted in binding buffer (50 mM Tris–HCl, pH 7.5, 2 mM β-mercaptoethanol, 3 mM EDTA, 100 mM NaCl, 0.2 µg/µl bovine serum albumin [BSA], 1 mM PMSF) for 1 hour at 4 °C with continuous rotation. The resin was washed 4 times with wash buffer containing 100 mM NaCl, proteins were eluted in wash buffer supplemented with 10 mM maltose and detected by western blotting^39^. As a negative control, HROB was incubated with the resin without the bait protein. For the experiment comparing phosphorylated and unphosphorylated variants of MCM8-9, pMCM8-9 (prepared as described in the purification section to preserve the phosphorylation status) was immobilized on amylose resin and was treated with λ-phosphatase for the unphosphorylated conditions, followed by three washes using the wash buffer, before incubation with wild type HROB.

### Structural Modeling

Sequences of human HROB (Q8N3J3), MCM8 (Q9UJA3) and MCM9 (Q9NXL9) were retrieved from UniProt database^67^ and used as input of mmseqs2 homology search program^68^ to generate a multiple sequence alignment (MSA) against the UniRef30 clustered database. Special care was taken in selecting only orthologs from the MSA of the proteins. For this, homologs of MCM8 and MCM9 sharing less than 35% sequence identity with their respective query and those of HROB sharing less than 25% were discarded. Homologs with less than 50% of coverage of the aligned region of their query were also discarded. In every MSA, in case several homologs belonged to the same species, only the one sharing highest sequence identity to the query was kept. Full-length sequences of the orthologs were retrieved and re-aligned with mafft^69^. To model the structure of the MCM8-MCM9 complex with and without HROB, the corresponding MSAs were concatenated. In the concatenated MSAs, when homologs of different subunits belonged to the same species they were aligned in a paired manner otherwise they were left unpaired. Final concatenated MSAs of HROB-MCM8-MCM9 contained 1247 sequences with 1888 positions and 222 paired sequences while that of MCM8-MCM9 contained 1247 sequences of 1539 positions with 690 matching sequences. The model of MCM8-MCM9 complex was calculated as a hexameric assembly using a stoichiometry of 3 for every subunit. Concatenated MSAs were used as inputs to generate 5 structural models for each of the HROB-MCM8-MCM9 trimer and of the MCM8-MCM9 hexamer using a local version of the ColabFold v1.5.2 interface^70^ running 3 iterations of the Alphafold2 v2.3.1 algorithm^71^ trained on the multimer dataset^72^ on a local HPC equipped with NVIDIA Ampere A100 80Go GPU cards. The five top models of each complexes converged toward very similar conformations and obtained good confidence and quality scores with pLDDTs in the range [73, 74.9] and [76.6, 80.1], pTMscore in the range [0.752, 0.773] and [0.706, 0.78] and ipTMscore in the range [0.705, 0.735] and [0.678, 0.757] for HROB-MCM8-MCM9 and MCM8-MCM9 hexamer, respectively. The models with the highest ipTMscores for both complexes were relaxed using rosetta relax protocols^73^ to remove steric clashes under constraints (std dev. of 2 Å for the interatomic distances) and were used for structural analysis. Molecular graphics and analyses were performed with UCSF ChimeraX^74^.

### Plasmids for cellular experiments

For immunoprecipitation, pMSCV-FLAG-HA-eGFP, -MCM8 and -MCM9 constructs were previously described^39^. To generate expression constructs for MCM8 (L387E) and MCM9 (M45E) mutants, Gateway entry vectors pDONR223-MCM8 and pDONR223-MCM9^39^ were subjected to site-directed mutagenesis by inverse PCR using primers listed in Supplementary Table S4. The mutant entry vectors were then recombined into pMSCV-FLAG-HA-DEST using LR clonase II (Thermo Fisher, 11791020).

### Immunoprecipitation

HEK293T were transiently transfected with pMSCV-FLAG-HA-MCM8, -MCM8 L387E, - MCM9, -MCM9 M45E or -GFP control (Transporter 5, Polysciences 26008-5) and harvested 3 days later for HA-immunoprecipitation as previously described^39^. Briefly, cell pellets were resuspended in mammalian cell lysis buffer (MCLB) (50 mM Tris-HCl pH 7.5, 1% IGEPAL CA-630) supplemented with 150 mM NaCl, protease inhibitor (Goldbio, GB-331) and phosphatase inhibitor cocktail (Sigma, 4906837001). Following incubation for 30 minutes at 4 °C, extracts were cleared by centrifugation and supernatants collected. The remaining pellets were subjected to a second round of extraction by resuspending them in MCLB supplemented with 500 mM NaCl and protease (Goldbio, GB-331) and phosphatase inhibitor cocktail (Sigma, 4906837001), and incubating them for an additional hour at 4 °C. After clearing these extracts by centrifugation, the salt concentration of the collected supernatants was adjusted to 150 mM NaCl with MCLB and combined with the supernatants from the first round of extraction. Combined lysates were then incubated with anti-HA agarose beads (Sigma, A2095) for 4 hours at 4 °C. After incubation, beads were washed four times in MCLB buffer supplemented with 150 mM NaCl and then boiled in LDS (lithium dodecyl sulfate) sample buffer (Thermo Fisher, NP0008) supplemented with beta-mercaptoethanol to elute the bound proteins.

### Phosphatase Assay

HEK293T cells from a near confluent 10 cm dish were harvested, pelleted by centrifugation, and resuspended in 500 µl lysis buffer (50 mM Tris-HCl pH 7.5, 150 mM NaCl, 1% IGEPAL CA-630) supplemented with protease inhibitor cocktail (Goldbio, GB-331). Following incubation on ice for 15 minutes, extracts were cleared by centrifugation. Phosphatase reactions (100 µl total volume) containing cleared lysates supplemented with MnCl_2_ (1 mM final concentration), 100 units lambda phosphatase (NEB, P0753S), and/or phosphatase inhibitor cocktail (2x final concentration) (Sigma, 4906837001) were prepared on ice and subsequently incubated at 30 °C for 1 hour. Reactions were then boiled with LDS sample buffer (Thermo Fisher, NP0008) supplemented with beta-mercaptoethanol, resolved by SDS-PAGE using a 6% acrylamide gel supplemented with 75 µM PhosBind reagent (APExBIO, F4002) and 150 µM MnCl_2_, and transferred to nitrocellulose membrane for immunoblotting.

### Antibodies

The following antibodies were used for immunoblotting: rabbit anti-HROB (Sigma HPA023393; 1:1000 dilution), rabbit anti-MCM8 (Proteintech 16451-1-AP; 1:5000 dilution), rabbit anti-MCM9 (EMD/Millipore ABE2603; 1:20000 dilution), mouse anti-HA (Sigma H3663; 1:2500 dilution), mouse anti-vinculin (Sigma V9131; 1:20000 dilution). For *in vitro* interaction studies, the following antibodies were used for western blotting: rabbit anti-MCM9 (Millipore ABE2603; 1:10,000 dilution) and mouse anti-FLAG (Sigma F1804; 1:1000 dilution) for FLAG-MCM8 and HROB-FLAG wild type and mutants.

### Flow-cell preparation for magnetic tweezers

In preparation for the assemblage of the flow-cell, two 60 mm x 24 mm cover slides (Menzel, ThermoScientific) were soaked subsequently in ultrapure water, acetone and isopropanol for 10 minutes each. Following each cleaning step, the slides were dried with N_2_ gas. The top slide was additionally modified with two holes of a diameter of ∼2 mm serving as an inlet and an outlet. The bottom slide was coated with a 1% w/vol polystyrene-toluene solution via spin-coating at 6000 rpm. To conclude the coating procedure, the bottom slide was placed in an oven for 90 minutes at 150 °C. The assembly of the flow-cell was finalized by placing a specifically cut out Parafilm layer in-between the top and the bottom slide, forming a cavity for the liquid flow. This arrangement was placed on a heating plate at 120 °C, thereby melting the Parafilm and permanently fusing the two slides together. In order to allow for specific surface binding, the assembled flow cell was incubated for 24 hours with a 50 µg/ml solution of anti-digoxigenin in phosphate-buffered saline (PBS).

Following this step, the anti-digoxigenin was replaced with a 20 mg/ml BSA solution in PBS and incubated for 24 hours. Subsequent to these two incubation steps, the flow-cell was installed inside the magnetic tweezer setup and connected to a pump. The flow-cell was first flushed with PBS to remove excess BSA. Then a solution of polystyrene beads with a size of 3.6 µm was flushed inside and left to incubate overnight, ensuring their firm attachment to the bottom slide of the flow-cell. These polystyrene beads will function as reference points for the subsequent magnetic-tweezer measurements. Prior to a magnetic tweezer measurement, 2.5 µl of streptavidin coated magnetic beads with a diameter of 1 µm (Dynabeads, MyOne, Invitrogen) were washed three times with PBS. The beads were then diluted in 2.5 µl PBS to which 1 µl of 40 ng/μl of Holliday junction construct was added. The mixture was left to incubate for 5 minutes at room temperature. Lastly, the incubated DNA-construct/magnetic bead mixture was resuspended in 100 µl PBS.

### HJ-containing DNA construct for magnetic tweezers

The Holliday junction construct was created from 4 parts: two linear DNA PCR fragments with lengths of 5326 bp (long fragment) and 4229 bp (short fragment), a biotin-modified DNA handle as well as a digoxigenin-modified DNA handle. The linear long and short fragments were synthesized via PCR from the plasmid pBluescript+1+2+4 (sequence available upon request) using the following primers: HJ-fragment 5326 bp Forward Primer, HJ-fragment 5326 bp Reverse Primer, HJ-fragment 4229 bp Forward Primer and HJ-fragment 4229 bp Reverse Primer. The fragments were subsequently digested with Nt.BbvCI and PspOMI (long fragment) or Nt.BbvCI and XhoI (short fragment), resulting in a mutually complementary 9 nt 3’-overhang of the two fragments. To ensure specific binding of the construct to the magnetic bead and flow-cell surface respectively, ∼550 bp biotin- and digoxigenin-modified DNA handles were produced by PCR from plasmid pBluescript II SK+ using biotin-modified dUTPs for the biotin handle and digoxigenin-modified dUTPs for digoxigenin handle^75^. The primers Handles Forward Primer and Handles Reverse Primer were used for both handles in pairs. Sequences of all used primers are listed in Supplementary Table S3. The biotin handle was digested with PspOMI and the digoxigenin handle with XhoI. The four distinct fragments (biotin-handle + long fragment + short fragment + digoxigenin handle) were then ligated in a single reaction, resulting in the 9546 bp Holliday junction construct (contour length ∼3.1 µm). Subsequently, the construct was gel-purified and stored at -20°C.

The overhang-construct^76, 77^ was prepared as follows: a 63 nt ssDNA gap was introduced into a 6.6 kbp dsDNA fragment using the nicking enzyme Nt.BbvCI. Thereafter, the gap was filled by hybridizing a 25 nt matching DNA oligomer sequence that exhibits an additional 40 nt polythimidine tail on its 5’-end. Subsequent ligation with biotin and digoxigenin modified handles yielded the final construct, exhibiting the overhang at ∼0.5 kbp distance from the surface attachment handle.

### Magnetic tweezers measurements on HJ substrate

The DNA-construct-magnetic bead mixture was flushed into the flow-cell and incubated for 150 seconds. Afterwards, the DNA molecules which did not specifically bind to the flow-cell surface were removed by washing with PBS. A suitable field of view, which included a stable reference bead and a magnetic bead tethered to the flow-cell surface by the DNA, was selected. To validate that the chosen construct was able to extrude the desired Holliday junction, it was examined if it was torsionally constrained. For that, 40 positive turns with respect to the helicity of the stretched dsDNA were applied at a force of 3.5 pN. Then, the force was lowered to 1 pN. If the result of this reduction in exerted force was a drastic change in the observable length of the DNA, the construct was confirmed as being torsionally constrained. When this prerequisite was met, 400 negative turns with respect to the helicity of the stretched dsDNA were applied at a force of 3.5 pN. The generated torque was sufficient to melt the hydrogen bonds of the DNA. Then the applied force was lowered to 1 pN in order to stimulate the formation of the Holliday junction. The incorporated mismatch in the center of the palindromic sequence served as an accessible starting point. After 1 hour, the formation of the junction could be observed as a shortening in DNA length. After successful formation of the Holliday junction, the total amount of negative turns was reduced to 50, in order to lower the torsional stiffness, thereby, making it easier for the investigated proteins to unwind the junction. PBS buffer was exchanged with the MCM buffer (25 mM HEPES-NaOH, 1 mM magnesium acetate, 1 mM DTT, 0.1 mg/ml BSA and 4 mM ATP) before measuring the proteins. Then, a force of 10 pN was applied, reducing the fluctuations of the magnetic bead to a minimum of around ± 9 nm. For each measurement, several baseline traces for the behavior of the junction in the MCM-buffer alone were recorded. The proteins were pre-mixed at the indicated concentrations and then flushed into the flow-cell, and the traces were recorded subsequently.

Single-molecule magnetic tweezer experiments with the overhang-construct were carried out in a custom-built magnetic tweezers setup at room temperature and 37 °C^78^. Flow cell preparation as well as first steps (pre-treatment, reference bead incubation, construct + magnetic bead incubation and subsequent flow cell incubation) are the same as described for the Holiday junction construct. Divergently, the measurements were conducted at 300 Hz using a real-time GPU accelerated image analysis^79^ allowing to investigate ∼10 molecules simultaneously. All measurements were performed at an average force of ∼15 pN (10-20 pN) in MCM-buffer (25 mM HEPES-NaOH pH 7.5, 1 mM magnesium acetate, 4 mM ATP, 1 mM DTT, 0.1 mg/ml BSA) whereas protein concentrations and molar ratios are as indicated in the figure.

## Data availability

The models are available in ModelArchive (modelarchive.org) with the accession codes ma-ji5l4 and ma-ma-zlpye for the MCM8-MCM9 hexamer and HROB-MCM8-MCM9 complex, respectively.

## Acknowledgements

We thank members of the Cejka Lab for their critical comments on this manuscript. We thank Boris Pfander (Max-Planck Institute of Biochemistry, Germany) for the CDK2-CycA expression construct and Roshan Singh Thakur (Protein facility, Institute for Research in Biomedicine, Bellinzona) for assistance with size exclusion chromatography. The Swiss National Science Foundation (SNSF) (Grants 310030_207588 and 310030_205199) and the European Research Council (ERC) (Grant 101018257) support the research in the Cejka laboratory. The French National Research Agency (ANR) (Grant ANR-21-CE44-0009-01) support the RG laboratory. The work in the Ciccia laboratory was supported by the NIH grant R01CA197774.

## Author contributions

A.A, R.G. and P.C conceived the idea. A.A designed and performed biochemical ensemble and mass photometry experiments. M.M. and M.G. performed the magnetic tweezer experiments under the supervision of R.S. H.B. and R.G. performed modeling experiments and mutation design. V.K performed the TEM experiments. JWH performed the experiments with cell extracts under the supervision of A.C. A.A and P.C wrote the manuscript. All authors contributed to prepare the final version of the manuscript.

## Supplementary Information

### Inventory

Extended Data Figures ED Fig.1 to 6

Extended Data Figure Legends

Supplementary Tables S1 to S4

### Extended Data Figure Legends

**ED Fig. 1.**
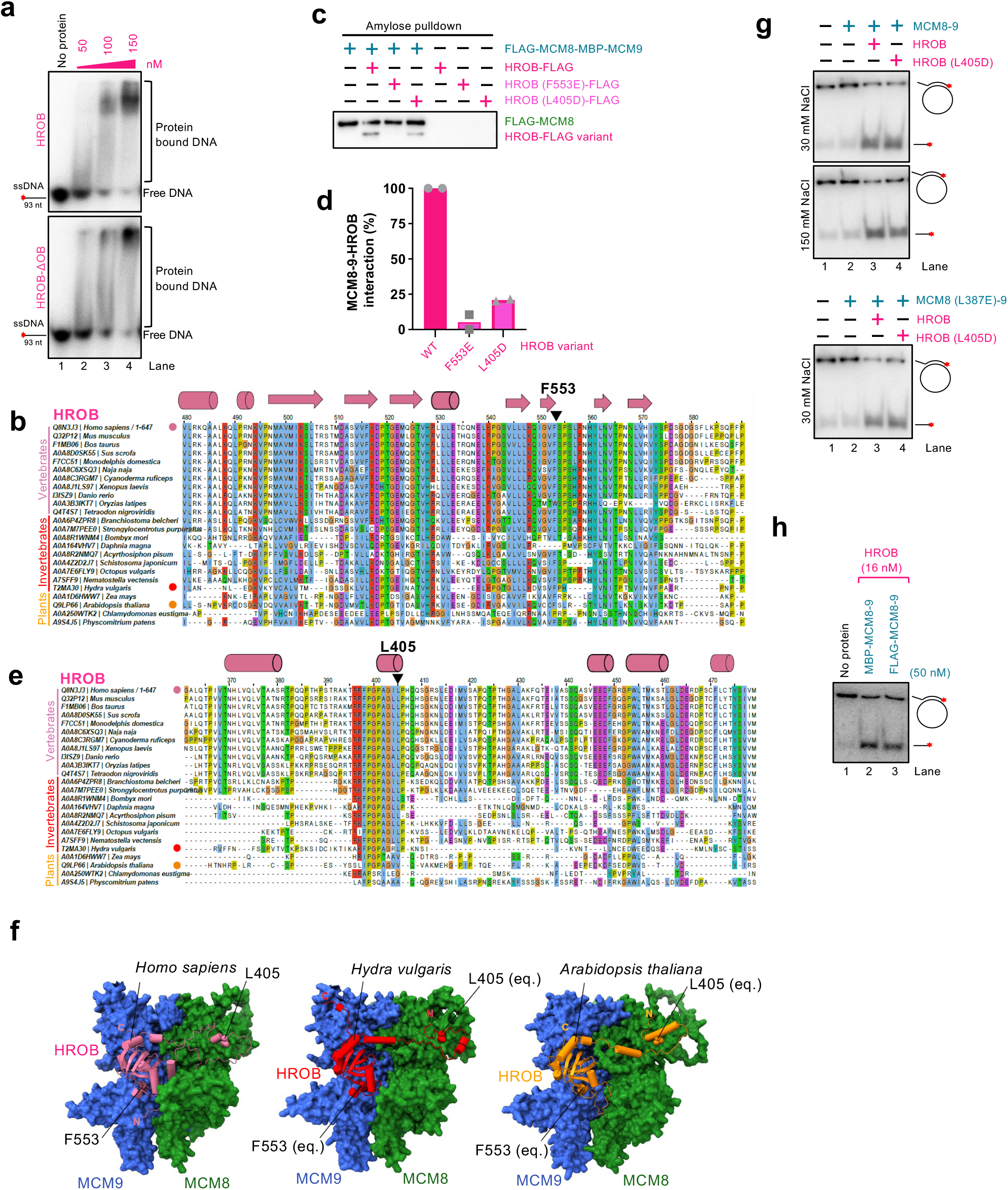
Interaction of HROB with MCM8-9. a. Electrophoretic mobility shift assay with human HROB or HROB-ΔOB and 93 nt-long ssDNA. The red asterisk indicates the position of the radioactive label. Shown is a representative experiment. b. Multiple sequence alignment of HROB showing evolutionary conservation of F553 and the surrounding region. c. MBP-tagged MCM8-9 was immobilized on amylose resin and incubated with HROB or variants at 100 mM NaCl. Bound proteins were eluted and visualized by western blotting with anti-FLAG antibody, detecting FLAG-tagged MCM8 and FLAG-tagged HROB. d. Quantification of assays such as shown in panel c; Data points show range; n = 2. The values were normalized to wild type HROB (lane 2). e. Multiple sequence alignment of HROB showing the conservation of L405 and the surrounding region. f. A cartoon showing a conservation of the interaction of HROB with MCM8-9 in various species, modelled by AlphaFold2. g. DNA unwinding with HROB (L405D) (25 nM) and MCM8-9 (50 nM) to test for the impact of disrupting HROB interaction with MCM8, carried out at indicated NaCl concentrations using circular M13-based ssDNA. The red asterisk indicates the position of the radioactive label. Shown is a representative experiment. h. A comparison of DNA unwinding by FLAG-MCM9 and untagged MCM8, or MBP-MCM9 and FLAG-MCM8, purified as a complex, in the presence of HROB using circular M13-based ssDNA. The red asterisk indicates the position of the radioactive label. The MBP and FLAG tags were not cleaved from the corresponding proteins. Shown is a representative experiment.

**ED Fig. 2.**
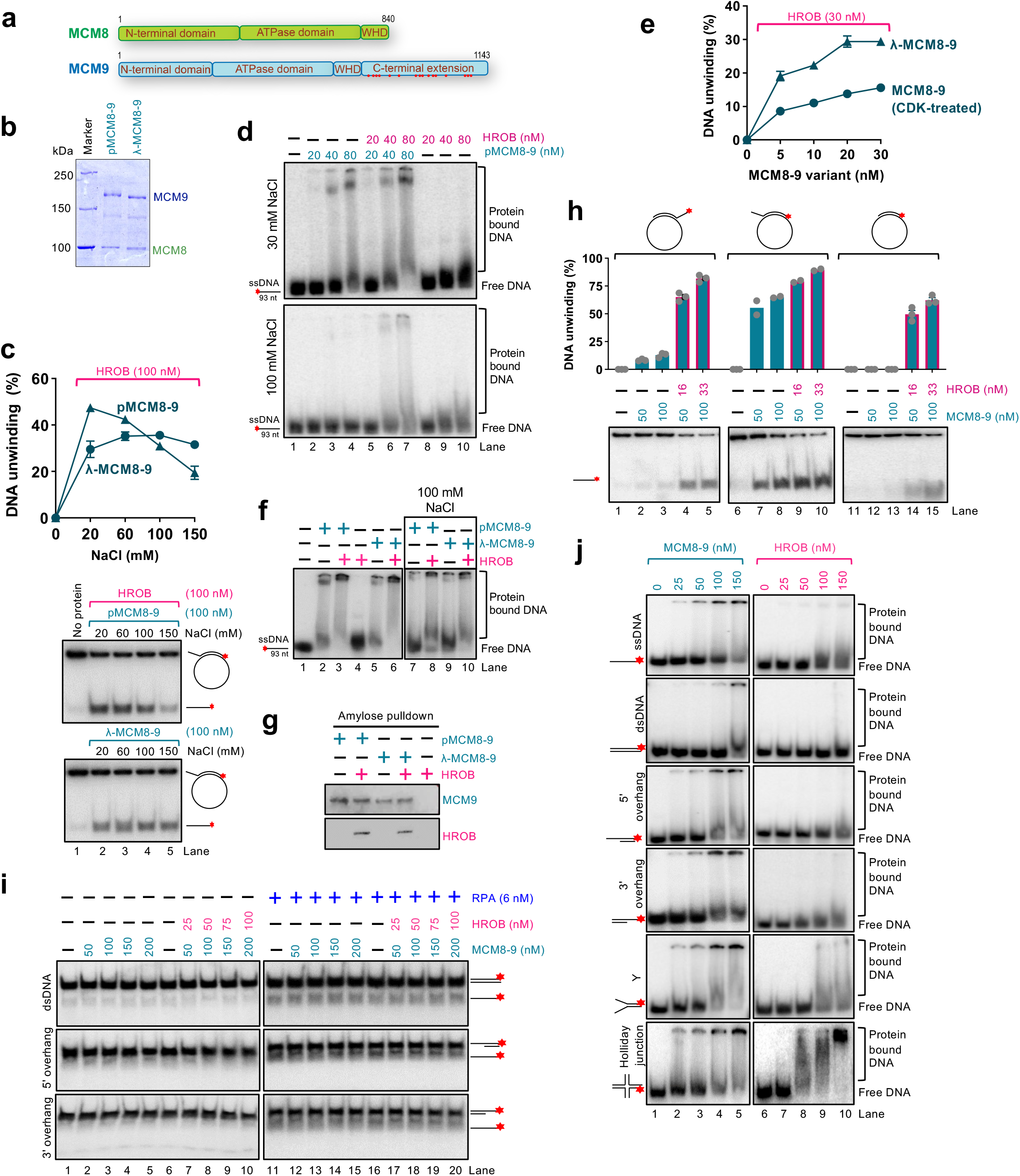
Regulation of MCM8-9 by phosphorylation and DNA binding and unwinding specificity of MCM8-9. a. A schematic representation of the main domains in human MCM8 and MCM9. WHD, winged-helix domain. The red dots represent approximate positions of 16 putative consensus CDK phosphorylation sites present in the unique C terminal tail of MCM9 (T673, S703, S707, S711, S762, S802, T879, S883, S890, S915, S942, S952, T977, T1064, S1073 and S1088). b. MCM8-9 heterodimer variants prepared with phosphatase inhibitors (phosphorylated MCM8-9, pMCM8-9), or without phosphatase inhibitors and treated with λ phosphatase during purification (dephosphorylated MCM8-9, λ-MCM8-9). The phosphorylated form of MCM9 shows a mobility shift in gel electrophoresis. c. DNA unwinding assays to compare the activities of phosphorylated (pMCM8-9) and dephosphorylated (λ-MCM8-9) MCM8-9 variants (both 100 nM), with HROB, and its dependence on NaCl concentration using circular M13-based ssDNA. The red asterisk indicates the position of the radioactive label. Top, quantification; error bars, SEM; n = 3; bottom, a representative experiment. d. Electrophoretic mobility shift assay with pMCM8-9 in the presence or absence of HROB using a 93 nt long ssDNA at no added and 100 mM NaCl with 3 mM EDTA. The red asterisk indicates the position of the radioactive label. Panel shows representative experiments. e. Quantification of DNA unwinding experiments by human MCM8-9, either phosphorylated with model CDK, CDK2-CycA, MCM8-9 (CDK), or dephosphorylated, λ-MCM8-9, at 150 mM NaCl using circular M13-based ssDNA. Error bars, SEM; n = 3. f. Electrophoretic mobility shift assays with pMCM8-9 or λ-MCM8-9 (both 100 nM), in the presence or absence of HROB (100 nM) with or without added salt (100 mM NaCl) and 93 nt-long ssDNA with 3 mM EDTA. Panel shows a representative experiment. g. Physical interaction assay of pMCM8-9 or λ-MCM8-9 and HROB at 100 mM NaCl. The MBP-tagged MCM8-9 variant was immobilized on amylose resin, HROB was subsequently added, bound proteins were eluted, and subjected to Western blotting. MCM9 was detected by anti-MCM9 and HROB by anti-FLAG antibodies. h. DNA unwinding by pMCM8-9 with and without HROB, using M13-based substrates with 5’, 3’ or no overhang (all substrates contained 37 bp of duplex DNA, overhang length 40 nt). Top, quantification; error bars, SEM; n = 3; data points in lanes 6-10 show range, n = 2; bottom, a representative experiment. i. DNA unwinding by pMCM8-9 without or with HROB and RPA, using oligonucleotide-based DNA substrates with 1 mM ATP, 5 mM magnesium acetate and 15 mM NaCl. Representative assays are shown. The red asterisk indicates the position of the radioactive label. j. Electrophoretic mobility shift assays with pMCM8-9 or HROB, using various oligonucleotide-based DNA substrates with 3 mM EDTA. Representative assays are shown. The red asterisk indicates the position of the radioactive label.

**ED Fig. 3.**
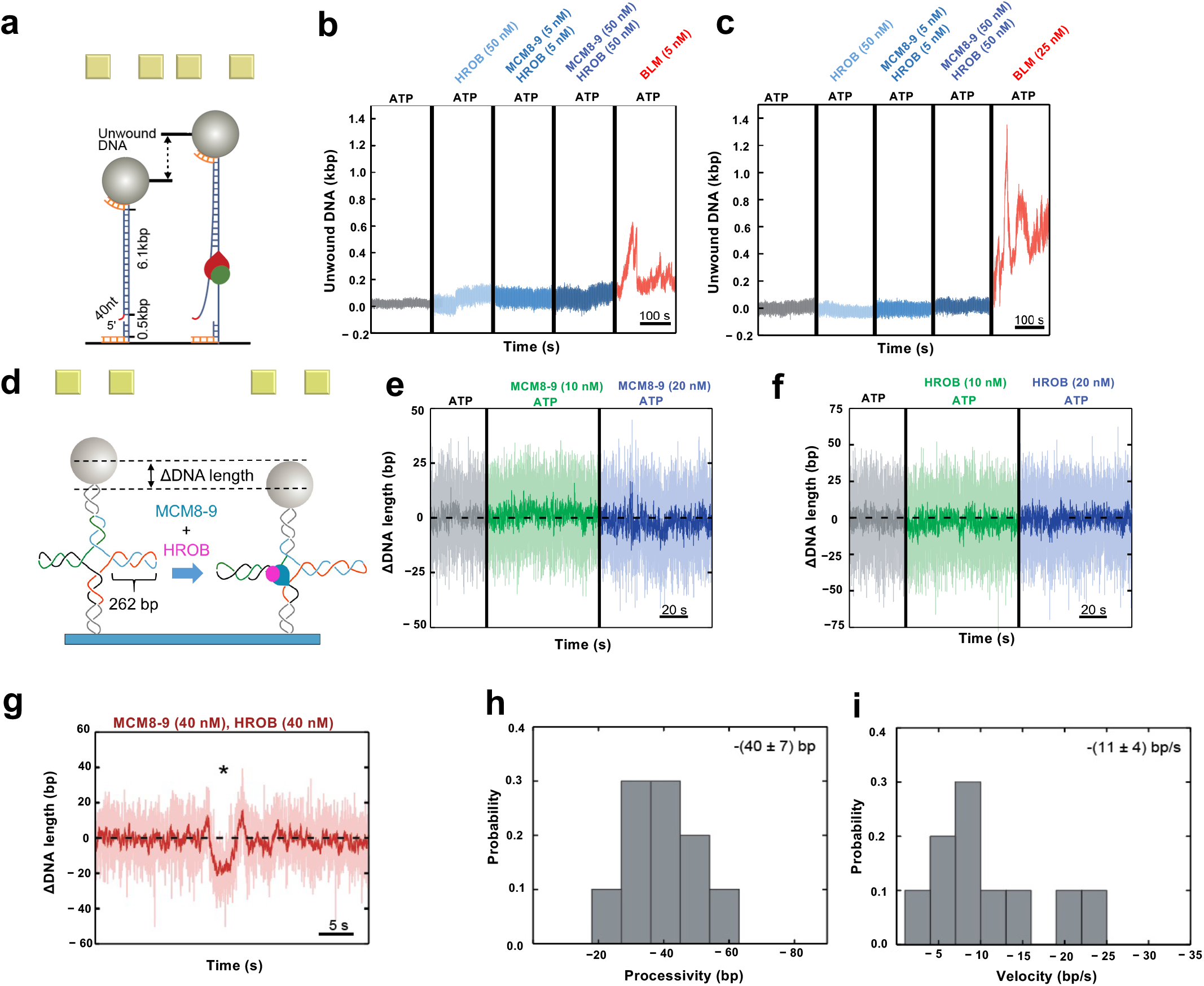
Magnetic tweezer experiments with the MCM8-9 helicase and HROB. a. A schematic representation of an overhanged DNA construct, the unwinding of which would lead to a change in bead position. b. Representative time trajectories of DNA length changes at room temperature under the following conditions: no protein (grey), 50 nM HROB (light blue), 5 nM MCM8-9:HROB (1:1) (blue) and 50 nM MCM8-9:HROB (1:1) (dark blue). Additionally, 5 nM BLM (red) was used as a positive control. Each measurement was performed at least 3 times. c. Experiments as in panel b, but at 37 °C. BLM concentration was 25 nM. d. A schematic representation of the magnetic tweezer setup and the investigated Holliday junction construct with 262 bp in each arm. When the protein ensemble is added, MCM8-9 with the cofactor HROB can translocate in the vertical direction, causing the arms of the Holliday junction to extent further, leading to a net-decrease in DNA length. e. Processing of the Holliday junction construct by MCM8-9 alone without HROB. No protein activity was detected. f. Processing of the Holliday junction construct by HROB alone. No protein activity was detected. g. An example trace of a net-downward event (highlighted by *) at a concentration of 40 nM for MCM8-9:HROB (1:1). a. Probability distribution of DNA unwinding processivity by MCM8-9 with HROB, with a mean of – (40 ± 7) bp, of events leading to DNA shortening, n = 10. b. Probability distribution of DNA unwinding velocity by MCM8-9 with HROB, with a mean of –(11 ± 4) bp sec^-1^, of events leading to DNA shortening, n = 10.

**ED Fig. 4.**
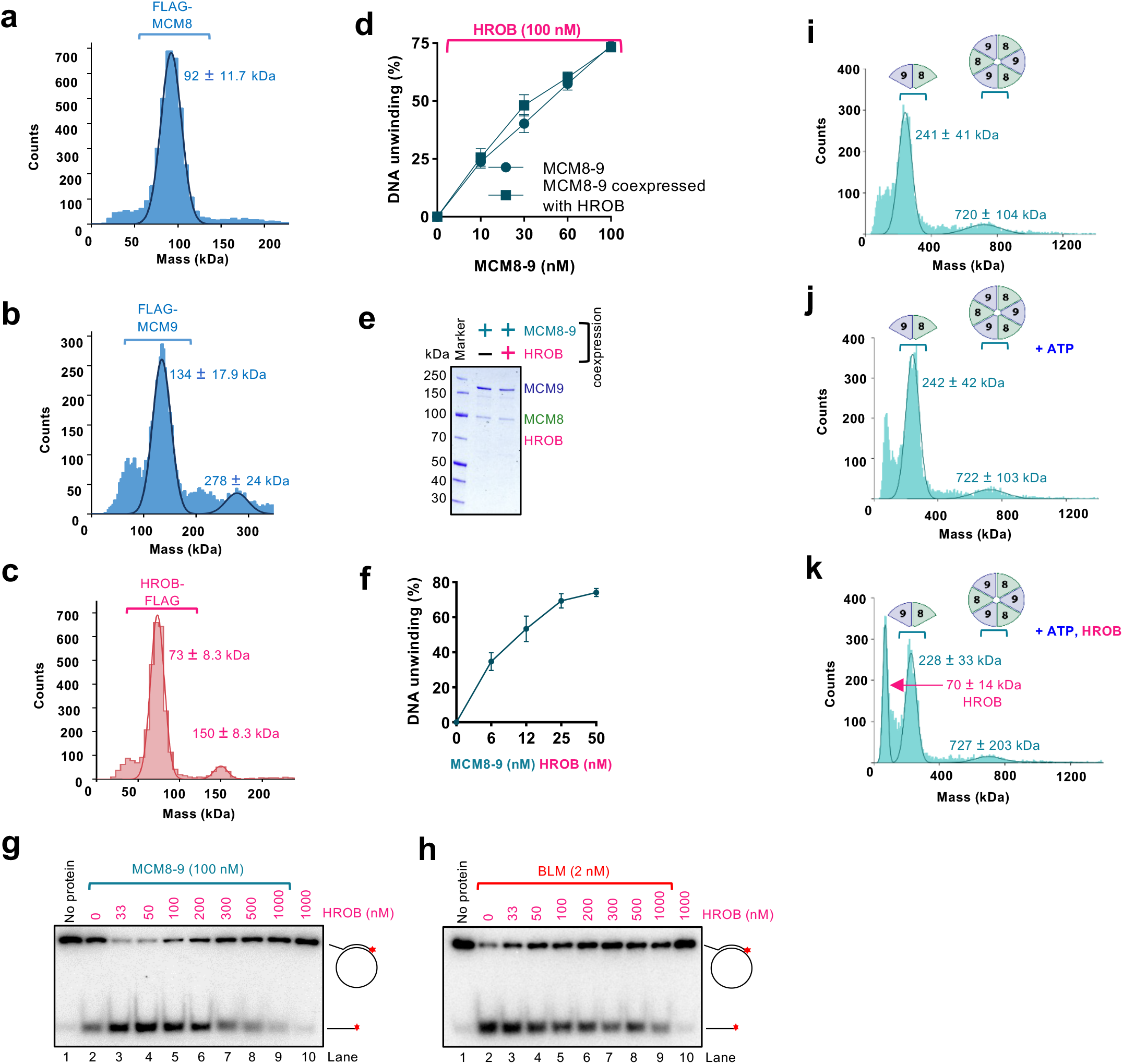
Analysis of MCM8-9 oligomerization. a. Measured molecular weight distribution of human FLAG-MCM8 (theoretical value 95 kDa including tag) using mass photometry. Error, SD. b. Measured molecular weight distribution of human FLAG-MCM9 (theoretical value 128 kDa including tag) using mass photometry. Error, SD. c. Measured molecular weight distribution of human HROB-FLAG (theoretical value 71 kDa including tag) using mass photometry. Error, SD. d. Helicase assays with MCM8-9 complex expressed as a heterodimer or co-expressed with HROB. MCM8-9 was 100 nM. As MCM8-9 did not co-purify with HROB, all reactions were supplemented with 100 nM HROB. Error bars, SEM; n = 3. e. Purified recombinant MCM8-9 complex expressed together with HROB, as used in panel d. HROB could not be co-purified with MCM8-9. f. Quantification of DNA unwinding assays using equimolar concentrations of MCM8-9 and HROB, as indicated, using M13-based circular ssDNA substrate. Error bars, SEM; n = 3. g. Representative DNA unwinding assays using human MCM8-9 in the presence of various HROB concentrations using M13-based circular ssDNA substrate. A representative experiment is shown. h. Representative DNA unwinding assays using human BLM in the presence of various HROB concentrations using M13-based circular ssDNA substrate. A representative experiment is shown. i. Measured molecular weight distribution of human FLAG-MCM8-MBP-MCM9 (MCM8-9, theoretical value 266 kDa for the heterodimer, 798 kDa for hexamer, including tags) using mass photometry. Errors, SD. j. Experiment as in panel i, but in a buffer additionally containing 2 mM ATP and 1 mM magnesium acetate. k. Experiment as in panel j, but together with HROB.

**ED Fig. 5.**
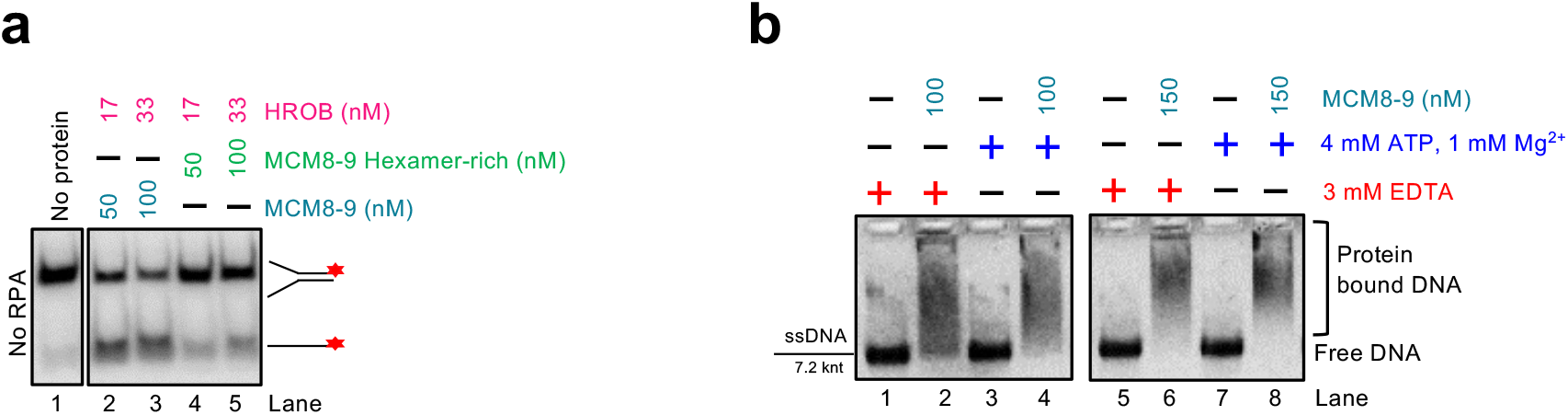
DNA binding and unwinding by MCM8-9. a. Representative DNA unwinding assays with standard and hexamer-rich MCM8-9 preparations in the presence HROB, using Y-structured DNA substrate. The red asterisk indicates the position of the radioactive label. b. Representative electrophoretic mobility shift assays with human MCM8-9 without or with ATP, using M13-based linear ssDNA (7.2 knt).

**ED Fig. 6.**
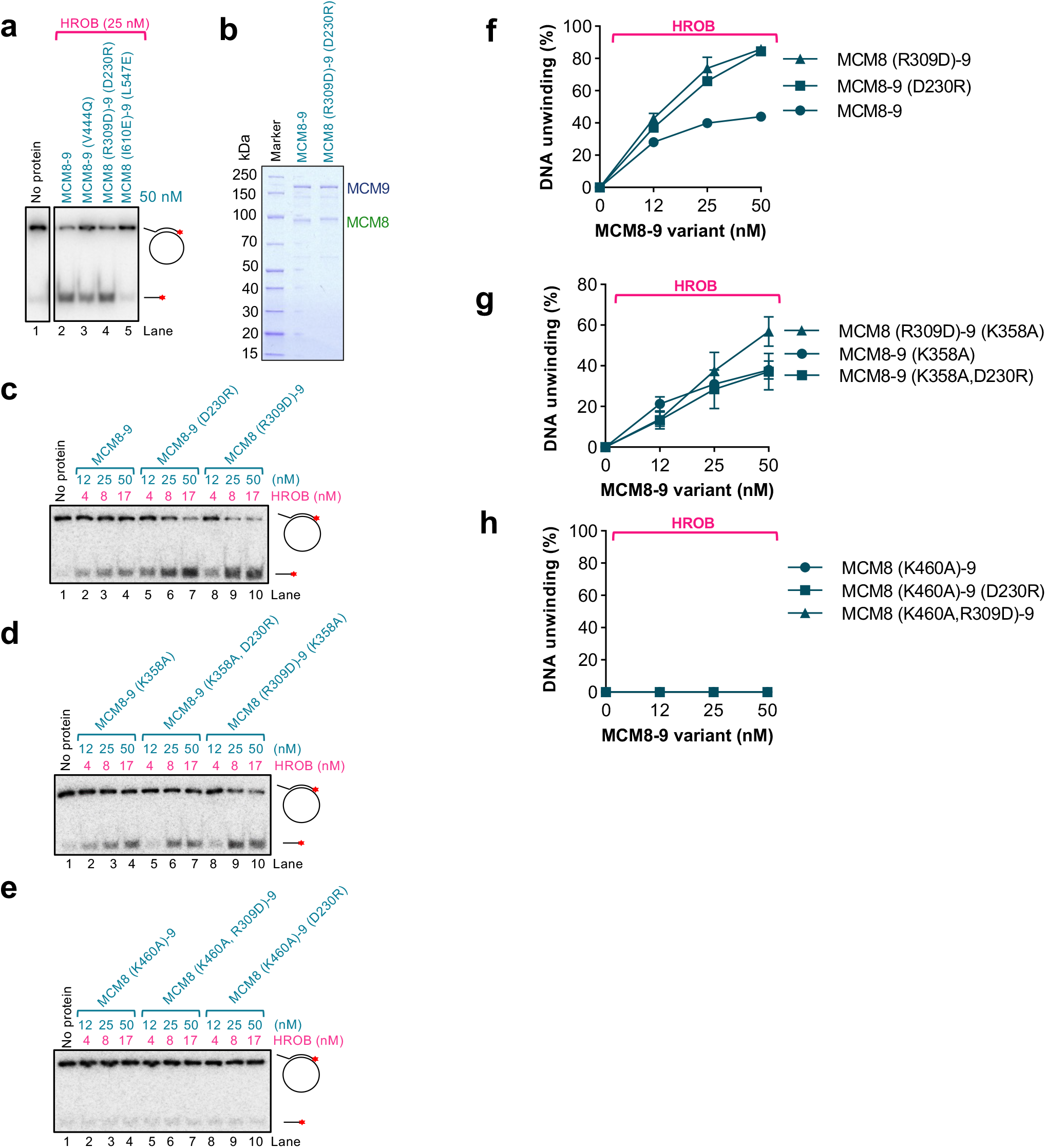
Analysis of MCM8-9 mutants disrupting the N-terminal part of interface II. a. Representative DNA unwinding assays with MCM8-9 variants (50 nM) mutated in interface II at 25 mM NaCl in the presence of HROB (25 nM) using M13-based circular ssDNA substrate. b. Wild type MCM8-9 and N-terminal interface II mutant MCM8 (R309D)-MCM9 (D230R) used in this study. c-e. MCM8-9 variants with disrupted N-terminal part of interface II, in combination with ATPase Walker A mutations, as indicated, at 25 mM NaCl in the presence of HROB at indicate concentrations using M13-based circular ssDNA substrate. Panel shows a representative experiment. f-h. Quantification of assays such as shown in panels c-e. HROB was used at 1/3 concentration of MCM8-9 (nM). Error bars, SEM; n = 3.

**Table S1:**
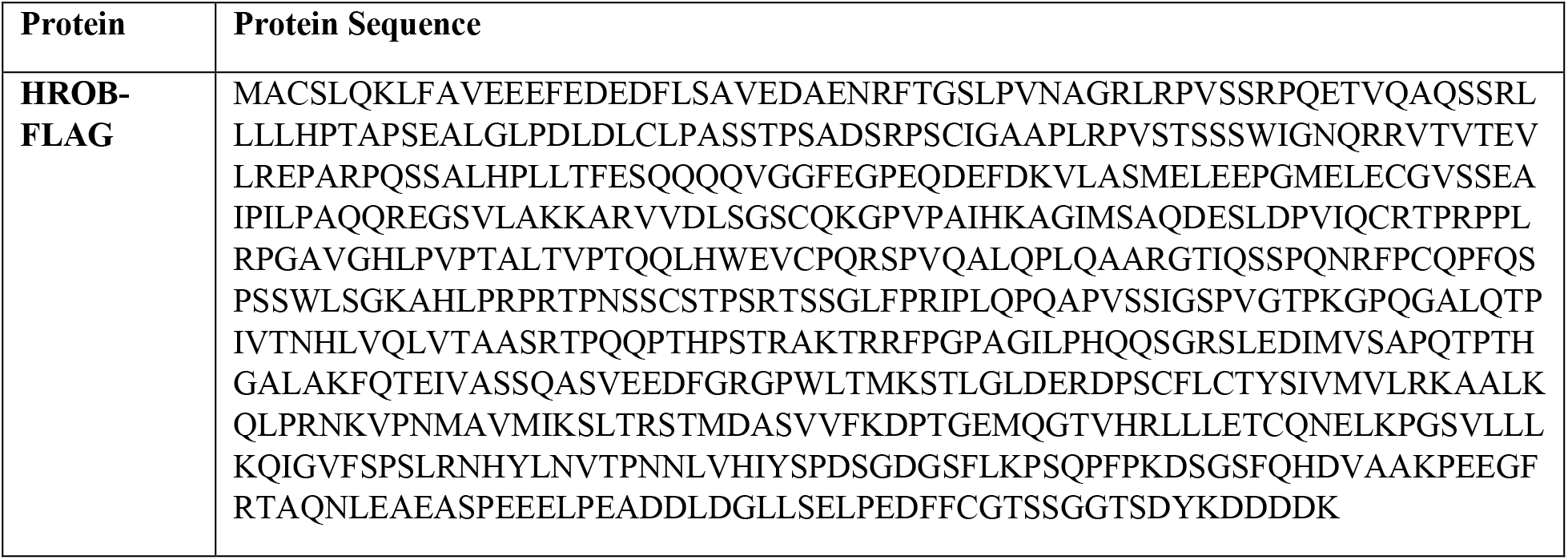
Protein sequence of HROB-FLAG used in this study.

**Table S2:**
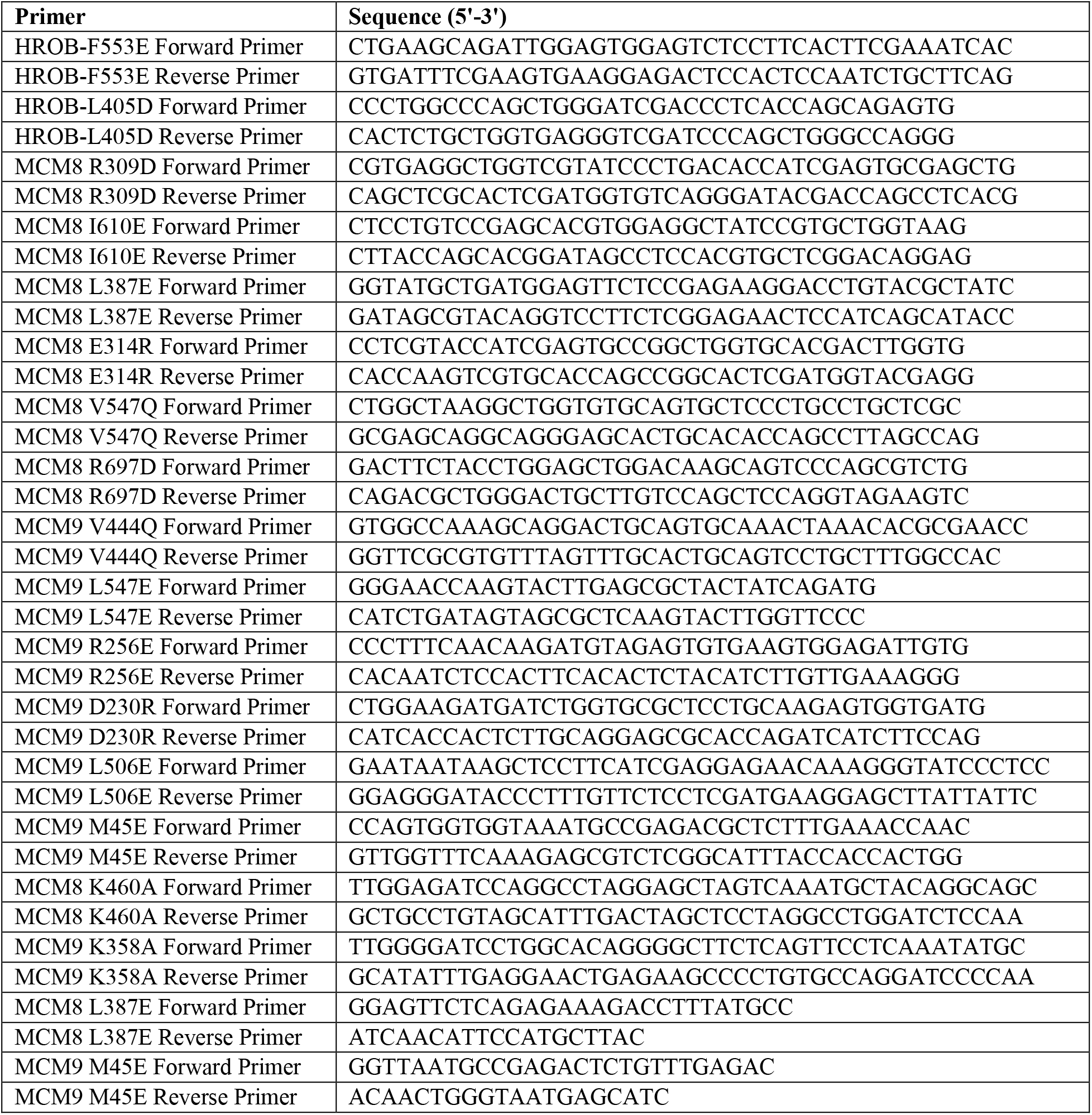
List of oligonucleotides used for site-directed mutagenesis to produce recombinant proteins.

**Table S3:**
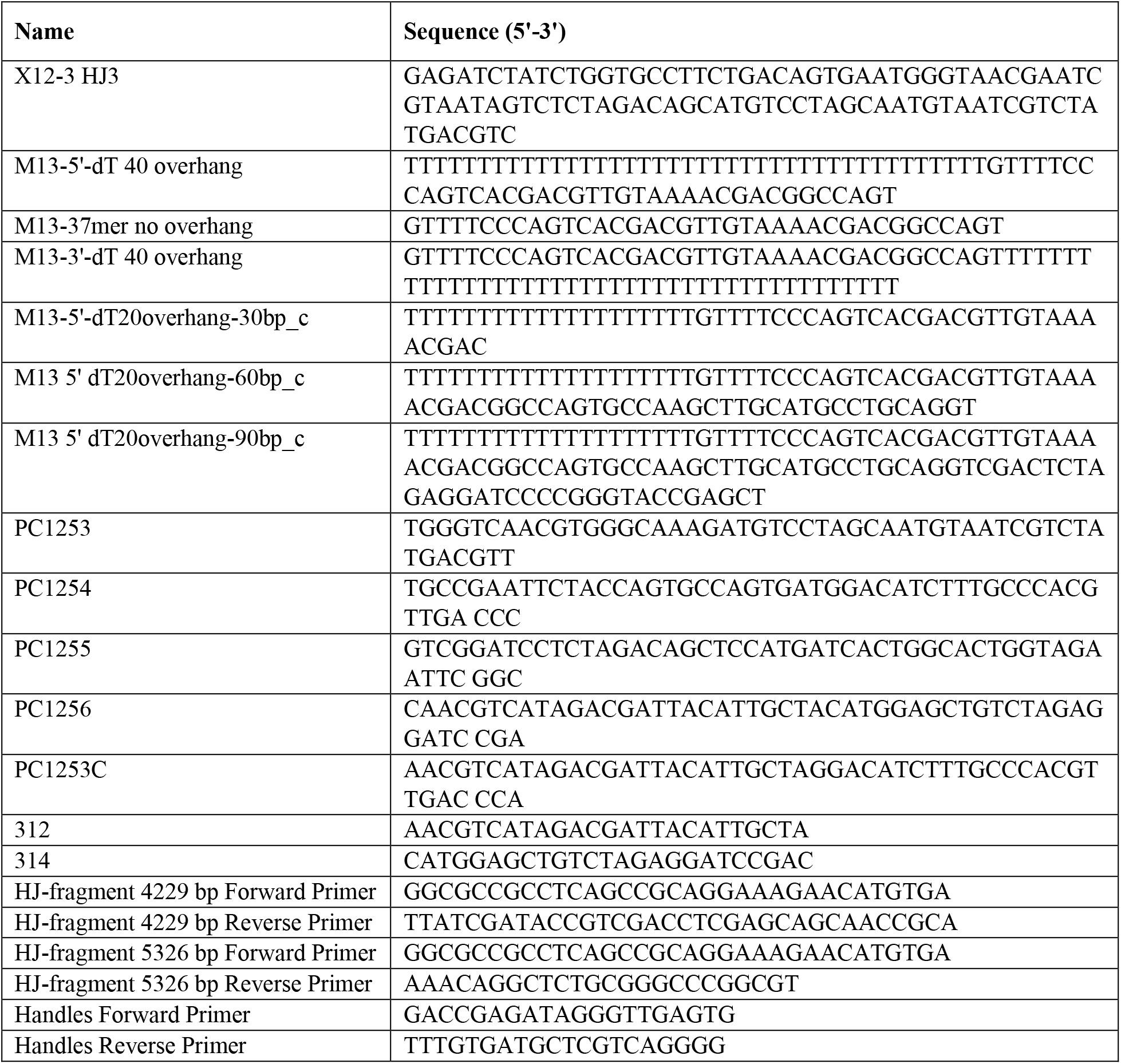
Sequence of oligonucleotides used for substrate preparation in this study.

**Table S4:**
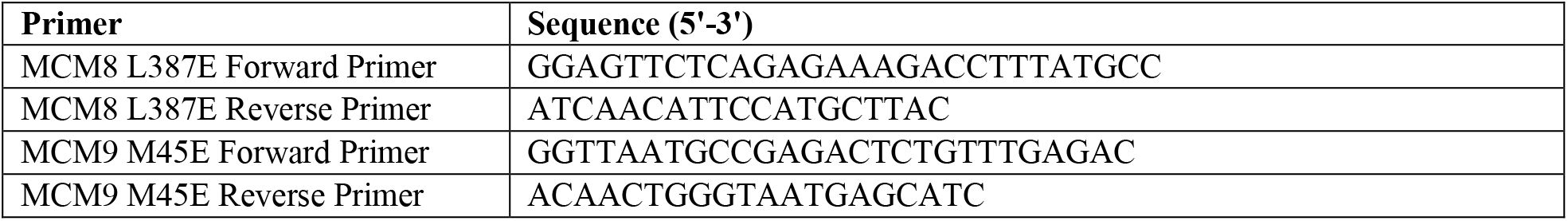
List of oligonucleotides used for site-directed mutagenesis to produce plasmids for cellular assays in this study.

## REFERENCES

1. Forsburg, S.L. Eukaryotic MCM proteins: beyond replication initiation. Microbiol Mol Biol Rev 68, 109–31 (2004).

2. Gao, Y. & Yang, W. Different mechanisms for translocation by monomeric and hexameric helicases. Curr Opin Struct Biol 61, 25–32 (2020).

3. Fernandez, A.J. & Berger, J.M. Mechanisms of hexameric helicases. Crit Rev Biochem Mol Biol 56, 621–639 (2021).

4. Maiorano, D., Lutzmann, M. & Mechali, M. MCM proteins and DNA replication. Curr Opin Cell Biol 18, 130–6 (2006).

5. Deegan, T.D. & Diffley, J.F. MCM: one ring to rule them all. Curr Opin Struct Biol 37, 145–51 (2016).

6. Coster, G. & Diffley, J.F.X. Bidirectional eukaryotic DNA replication is established by quasi-symmetrical helicase loading. Science 357, 314–318 (2017).

7. Douglas, M.E., Ali, F.A., Costa, A. & Diffley, J.F.X. The mechanism of eukaryotic CMG helicase activation. Nature 555, 265–268 (2018).

8. Miller, T.C.R., Locke, J., Greiwe, J.F., Diffley, J.F.X. & Costa, A. Mechanism of head-to-head MCM double-hexamer formation revealed by cryo-EM. Nature 575, 704–710 (2019).

9. Costa, A. et al. DNA binding polarity, dimerization, and ATPase ring remodeling in the CMG helicase of the eukaryotic replisome. Elife 3, e03273 (2014).

10. Lewis, J.S. et al. Mechanism of replication origin melting nucleated by CMG helicase assembly. Nature 606, 1007–1014 (2022).

11. Kelman, L.M., O’Dell, W.B. & Kelman, Z. Unwinding 20 Years of the Archaeal Minichromosome Maintenance Helicase. J Bacteriol 202(2020).

12. Schwacha, A. & Bell, S.P. Interactions between two catalytically distinct MCM subgroups are essential for coordinated ATP hydrolysis and DNA replication. Mol Cell 8, 1093–104 (2001).

13. Erzberger, J.P. & Berger, J.M. Evolutionary relationships and structural mechanisms of AAA+ proteins. Annu Rev Biophys Biomol Struct 35, 93–114 (2006).

14. Bochman, M.L., Bell, S.P. & Schwacha, A. Subunit organization of Mcm2-7 and the unequal role of active sites in ATP hydrolysis and viability. Mol Cell Biol 28, 5865–73 (2008).

15. Liu, Y., Richards, T.A. & Aves, S.J. Ancient diversification of eukaryotic MCM DNA replication proteins. BMC Evol Biol 9, 60 (2009).

16. Blanton, H.L., et al. REC, Drosophila MCM8, drives formation of meiotic crossovers. PLoS Genet 1, e40 (2005).

17. Kohl, K.P., Jones, C.D. & Sekelsky, J. Evolution of an MCM complex in flies that promotes meiotic crossovers by blocking BLM helicase. Science 338, 1363–5 (2012).

18. Crismani, W. et al. MCM8 is required for a pathway of meiotic double-strand break repair independent of DMC1 in Arabidopsis thaliana. PLoS Genet 9, e1003165 (2013).

19. Hutchins, J.R. et al. Proteomic data on the nuclear interactome of human MCM9. Data Brief 6, 410–5 (2016).

20. Li, J. et al. Structural study of the N-terminal domain of human MCM8/9 complex. Structure 29, 1171–1181 e4 (2021).

21. Weng, Z. et al. Structural and mechanistic insights into the MCM8/9 helicase complex. eLife (2023).

22. McKinzey, D.R. et al. Motifs of the C-terminal domain of MCM9 direct localization to sites of mitomycin-C damage for RAD51 recruitment. J Biol Chem 296, 100355 (2021).

23. Dou, X. et al. Minichromosome maintenance complex component 8 mutations cause primary ovarian insufficiency. Fertil Steril 106, 1485–1489 e2 (2016).

24. Jin, H. et al. Identification of potential causal variants for premature ovarian failure by whole exome sequencing. BMC Med Genomics 13, 159 (2020).

25. Biswas, L. et al. Meiosis interrupted: the genetics of female infertility via meiotic failure. Reproduction 161, R13–R35 (2021).

26. Tucker, E.J. et al. Meiotic genes in premature ovarian insufficiency: variants in HROB and REC8 as likely genetic causes. Eur J Hum Genet 30, 219–228 (2022).

27. Lutzmann, M. et al. MCM8- and MCM9 Deficiencies Cause Lifelong Increased Hematopoietic DNA Damage Driving p53-Dependent Myeloid Tumors. Cell Rep 28, 2851–2865 e4 (2019).

28. Hao, J. et al. Downregulation of MCM8 expression restrains the malignant progression of cholangiocarcinoma. Oncol Rep 46(2021).

29. Soares de Lima, Y., et al. Germline and Somatic Whole-Exome Sequencing Identifies New Candidate Genes Involved in Familial Predisposition to Serrated Polyposis Syndrome. Cancers (Basel) 13(2021).

30. Wang, X. et al. MCM8 is regulated by EGFR signaling and promotes the growth of glioma stem cells through its interaction with DNA-replication-initiating factors. Oncogene 40, 4615–4624 (2021).

31. Yang, S. et al. MCM4 is a novel prognostic biomarker and promotes cancer cell growth in glioma. Front Oncol 12, 1004324 (2022).

32. Lutzmann, M. et al. MCM8- and MCM9-deficient mice reveal gametogenesis defects and genome instability due to impaired homologous recombination. Mol Cell 47, 523–34 (2012).

33. Nishimura, K. et al. Mcm8 and Mcm9 form a complex that functions in homologous recombination repair induced by DNA interstrand crosslinks. Mol Cell 47, 511–22 (2012).

34. Hartmann, M., Kohl, K.P., Sekelsky, J. & Hatkevich, T. Meiotic MCM Proteins Promote and Inhibit Crossovers During Meiotic Recombination. Genetics 212, 461–468 (2019).

35. Liu, K. et al. Aberrantly expressed HORMAD1 disrupts nuclear localization of MCM8-MCM9 complex and compromises DNA mismatch repair in cancer cells. Cell Death Dis 11, 519 (2020).

36. Griffin, W.C. et al. A multi-functional role for the MCM8/9 helicase complex in maintaining fork integrity during replication stress. Nat Commun 13, 5090 (2022).

37. Hustedt, N. et al. Control of homologous recombination by the HROB-MCM8-MCM9 pathway. Genes Dev 33, 1397–1415 (2019).

38. Saredi, G. & Rouse, J. Ways to unwind with HROB, a new player in homologous recombination. Genes Dev 33, 1293–1294 (2019).

39. Huang, J.W. et al. MCM8IP activates the MCM8-9 helicase to promote DNA synthesis and homologous recombination upon DNA damage. Nat Commun 11, 2948 (2020).

40. Lee, K.Y. et al. MCM8-9 complex promotes resection of double-strand break ends by MRE11-RAD50-NBS1 complex. Nature communications 6, 7744 (2015).

41. Park, J. et al. The MCM8-MCM9 complex promotes RAD51 recruitment at DNA damage sites to facilitate homologous recombination. Mol Cell Biol 33, 1632–44 (2013).

42. Traver, S. et al. MCM9 Is Required for Mammalian DNA Mismatch Repair. Mol Cell 59, 831–9 (2015).

43. Liu, Q. et al. Pathogenic germline MCM9 variants are rare in Australian Lynch-like syndrome patients. Cancer Genet 209, 497–500 (2016).

44. Golubicki, M. et al. Germline biallelic Mcm8 variants are associated with early-onset Lynch-like syndrome. JCI Insight 5(2020).

45. Gambus, A. & Blow, J.J. Mcm8 and Mcm9 form a dimeric complex in Xenopus laevis egg extract that is not essential for DNA replication initiation. Cell Cycle 12, 1225–32 (2013).

46. Natsume, T. et al. Acute inactivation of the replicative helicase in human cells triggers MCM8-9-dependent DNA synthesis. Genes Dev 31, 816–829 (2017).

47. Wang, C. et al. C17orf53 is identified as a novel gene involved in inter-strand crosslink repair. DNA Repair (Amst) 95, 102946 (2020).

48. Piovesan, D. et al. MobiDB: 10 years of intrinsically disordered proteins. Nucleic Acids Res 51, D438–D444 (2023).

49. Ishimi, Y. Regulation of MCM2-7 function. Genes Genet Syst 93, 125–133 (2018).

50. Ishimi, Y., Komamura-Kohno, Y., You, Z., Omori, A. & Kitagawa, M. Inhibition of Mcm4,6,7 helicase activity by phosphorylation with cyclin A/Cdk2. J Biol Chem 275, 16235–41 (2000).

51. Bochman, M.L. & Schwacha, A. The Mcm complex: unwinding the mechanism of a replicative helicase. Microbiol Mol Biol Rev 73, 652–83 (2009).

52. Rothenberg, E., Trakselis, M.A., Bell, S.D. & Ha, T. MCM forked substrate specificity involves dynamic interaction with the 5’-tail. J Biol Chem 282, 34229–34 (2007).

53. Moreau, M.J., McGeoch, A.T., Lowe, A.R., Itzhaki, L.S. & Bell, S.D. ATPase site architecture and helicase mechanism of an archaeal MCM. Mol Cell 28, 304–14 (2007).

54. Kasaciunaite, K. et al. Competing interaction partners modulate the activity of Sgs1 helicase during DNA end resection. EMBO J 38, e101516 (2019).

55. Ceppi, I. et al. CtIP promotes the motor activity of DNA2 to accelerate long-range DNA end resection. Proc Natl Acad Sci U S A 117, 8859–8869 (2020).

56. Wasserman, M.R., Schauer, G.D., O’Donnell, M.E. & Liu, S. Replication Fork Activation Is Enabled by a Single-Stranded DNA Gate in CMG Helicase. Cell 178, 600–611 e16 (2019).

57. Kose, H.B., Xie, S., Cameron, G., Strycharska, M.S. & Yardimci, H. Duplex DNA engagement and RPA oppositely regulate the DNA-unwinding rate of CMG helicase. Nat Commun 11, 3713 (2020).

58. Li, N. et al. Structure of the eukaryotic MCM complex at 3.8 A. Nature 524, 186–91 (2015).

59. Bochman, M.L. & Schwacha, A. The Mcm2-7 complex has in vitro helicase activity. Mol Cell 31, 287–93 (2008).

60. Meagher, M., Epling, L.B. & Enemark, E.J. DNA translocation mechanism of the MCM complex and implications for replication initiation. Nat Commun 10, 3117 (2019).

61. Langston, L.D. et al. Mcm10 promotes rapid isomerization of CMG-DNA for replisome bypass of lagging strand DNA blocks. Elife 6(2017).

62. Looke, M., Maloney, M.F. & Bell, S.P. Mcm10 regulates DNA replication elongation by stimulating the CMG replicative helicase. Genes Dev 31, 291–305 (2017).

63. Balbo Pogliano, C., et al. The CDK1-TOPBP1-PLK1 axis regulates the Bloom’s syndrome helicase BLM to suppress crossover recombination in somatic cells. Sci Adv 8, eabk0221 (2022).

64. Anand, R., Pinto, C. & Cejka, P. Methods to Study DNA End Resection I: Recombinant Protein Purification. Methods Enzymol 600, 25–66 (2018).

65. Acharya, A. et al. Distinct RPA domains promote recruitment and the helicase-nuclease activities of Dna2. Nat Commun 12, 6521 (2021).

66. Cannavo, E. et al. Regulation of the MLH1-MLH3 endonuclease in meiosis. Nature 586, 618–622 (2020).

67. UniProt, C. UniProt: the universal protein knowledgebase in 2021. Nucleic Acids Res 49, D480–D489 (2021).

68. Steinegger, M. & Soding, J. MMseqs2 enables sensitive protein sequence searching for the analysis of massive data sets. Nat Biotechnol 35, 1026–1028 (2017).

69. Katoh, K. & Standley, D.M. MAFFT multiple sequence alignment software version 7: improvements in performance and usability. Mol Biol Evol 30, 772–80 (2013).

70. Mirdita, M. et al. ColabFold: making protein folding accessible to all. Nat Methods 19, 679–682 (2022).

71. Jumper, J. et al. Highly accurate protein structure prediction with AlphaFold. Nature 596, 583–589 (2021).

72. Evans, R., et al. Protein complex prediction with AlphaFold-Multimer. bioRxiv, 2021. 10.04.463034 (2022).

73. Leman, J.K. et al. Macromolecular modeling and design in Rosetta: recent methods and frameworks. Nat Methods 17, 665–680 (2020).

74. Pettersen, E.F. et al. UCSF ChimeraX: Structure visualization for researchers, educators, and developers. Protein Sci 30, 70–82 (2021).

75. Luzzietti, N., Knappe, S., Richter, I. & Seidel, R. Nicking enzyme-based internal labeling of DNA at multiple loci. Nat Protoc 7, 643–53 (2012).

76. Levikova, M., Klaue, D., Seidel, R. & Cejka, P. Nuclease activity of Saccharomyces cerevisiae Dna2 inhibits its potent DNA helicase activity. Proc Natl Acad Sci U S A 110, E1992–2001 (2013).

77. Grigaitis, R. et al. Phosphorylation of the RecQ Helicase Sgs1/BLM Controls Its DNA Unwinding Activity during Meiosis and Mitosis. Dev Cell 53, 706–723 e5 (2020).

78. Klaue, D. & Seidel, R. Torsional stiffness of single superparamagnetic microspheres in an external magnetic field. Phys Rev Lett 102, 028302 (2009).

79. Huhle, A. et al. Camera-based three-dimensional real-time particle tracking at kHz rates and Angstrom accuracy. Nat Commun 6, 5885 (2015).

80. Vilar, E. et al. Microsatellite instability due to hMLH1 deficiency is associated with increased cytotoxicity to irinotecan in human colorectal cancer cell lines. Br J Cancer 99, 1607–12 (2008).

81. Mandal, R. et al. Genetic diversity of tumors with mismatch repair deficiency influences anti-PD-1 immunotherapy response. Science 364, 485–491 (2019).

